# Mucin promotes pneumococcal colonization via autolysis suppression and tryptophan-mediated immune modulation

**DOI:** 10.64898/2026.11.06.764952

**Authors:** Cydney N. Johnson, Matthew W. Frank, Yuri Laguna-Terai, William MacCain, Tyler S. Simmons, Svetlana Zamakhaeva, Natalia Korotkova, Carol Aherne, Elaine Tuomanen, Christopher M. Evans, Katharina Ribbeck, Jason W. Rosch

## Abstract

*Streptococcus pneumoniae* is a leading human respiratory pathogen, yet its success depends not on disease but on its ability to persist as an asymptomatic colonizer of the nasopharynx. The nasopharyngeal mucosa is lined with mucus, whose heavily O-glycosylated mucin proteins are the most abundant macromolecules in the pneumococcal natural niche. While mucus is canonically viewed as a host defense barrier, we found that mucin actively reprograms pneumococcal physiology to promote colonization through two distinct mechanisms. First, mucin suppresses pneumococcal autolysis by specifically inhibiting the enzymatic activity of the major autolysin LytA, prolonging bacterial survival and restricting release of the cholesterol-dependent cytotoxin pneumolysin, thereby reducing host cell damage. Second, mucin induces *de novo* biosynthesis and extracellular release of tryptophan, an essential amino acid that humans cannot synthesize and that serves as a substrate for host immunomodulatory pathways. Tryptophan biosynthesis was critical for pneumococcal survival against macrophage killing, and its disruption abolished both colonization and invasive disease. Transcriptomic analysis revealed that tryptophan biosynthesis enables the pneumococcus to survive a bactericidal macrophage state while preventing macrophage transition to proliferative tissue repair. Together, these findings reveal that mucin is not merely a barrier but an active signal that reprograms the pneumococcus toward a persistent, host-tolerant state, identifying a distinctive repertoire of colonization mechanisms that allow a virulent respiratory pathogen to sustain long-term asymptomatic carriage in its human reservoir.

## INTRODUCTION

*Streptococcus pneumoniae* (the pneumococcus) is a remarkably adaptable human pathobiont, playing dual roles of asymptomatic carriage in the nasopharynx and invasive disease in the lower respiratory tract and bloodstream. The pneumococcus is the most common cause of community-acquired pneumonia in those under five and over sixty-five years old^1^, and is a leading cause of death in children. Even in the era of antibiotics, community-acquired pneumonia causes 0.6 to 1.2 million annual deaths despite being considered one of the most vaccine-preventable diseases^2,3^. What sets the pneumococcus apart from most notorious respiratory pathogens is not its capacity for disease, but rather its ability to persist. The human nasopharynx is the only natural reservoir for *S. pneumoniae*, and the bacterium is carried in the nasopharynx of 40-95% of infants and 1-10% of adults at any given time^4^. Colonization is the prerequisite for disease development and transmission^5^, yet the mechanisms that allow this bacterium to maintain in the human reservoir remain poorly understood. Understanding this part of the pneumococcal lifecycle will reveal fundamental biology behind the pneumococci’s success as a pathogen.

The nasal mucosa is lined with mucus, a viscoelastic hydrogel that serves as a protective barrier between the host epithelium and the environment^6^. The most abundant macromolecule of mucus are mucin proteins; the upper respiratory tract is dominated by the gel-forming mucins MUC5AC and MUC5B, with MUC2 at low levels^7,8^. Mucus is canonically thought of as an innate defense, trapping and clearing pathogens and debris^6^. The pneumococcus survives at the respiratory mucosal sites and is in constant contact with mucus. The pneumococcus has known strategies to contend with mucus, including neuraminidase-mediated adherence, capsule-mediated evasion of entrapment, and pilus-driven agglutination^9–12^. Yet these discoveries frame mucus primarily as an obstacle that the pneumococcus must overcome. An alternative and largely unexplored possibility is that mucus, and its components, can actively signal to colonizing bacteria, reshaping bacterial physiology in a manner that promotes persistence rather than invasion. In the intestinal tract, mucin serves as both nutrient source and environmental cue for the resident microbiota. However, whether mucin plays a similar role for respiratory colonizers such as the pneumococcus is unknown.

A clue to how mucin might actively promote colonization comes from tryptophan. Metabolomic profiling of the mouse nasopharynx during pneumococcal colonization revealed that tryptophan is one of only a handful of metabolites significantly elevated in the upper airway during pneumococcal carriage^13^, yet the source of this increase was not determined. Tryptophan is an essential amino acid that humans cannot synthesize and must obtain from their diet or bacteria that possess the biosynthetic pathway. Beyond protein synthesis, tryptophan occupies a unique position at the host-pathogen interface. Macrophages catabolize tryptophan via indoleamine 2,3-dioxygenase (IDO) as part of the kynurenine pathway; this depletion serves as an antibacterial strategy that starves auxotrophic bacteria in the phagolysosome^14,15^. This same pathway leads to the production of immunomodulatory kynurenine and indole that bind to the aryl hydrocarbon receptor (AhR) and drive macrophage polarization towards anti-inflammatory states^16^. In the gut, microbial tryptophan metabolism through these pathways is a cornerstone of mucosal homeostasis^17–19^. Several genetic screens have implicated the pneumococcal tryptophan biosynthetic locus at the host-pathogen interface^20–22^, but whether the pneumococcus itself synthesizes and releases tryptophan to actively modulate host immunity, and whether this is regulated by mucin, has not been tested.

Here we show that mucin induces two previously unrecognized changes in the pneumococcus that are directly implicated in successful colonization of the human reservoir. First, mucin suppresses pneumococcal autolysis by specifically inhibiting cell wall amidase activity, restricting release of the pore-forming toxin pneumolysin. This reduces host cell damage while prolonging bacterial survival, both essential for population maintenance in the nasopharynx. Second, mucin triggers *de novo* biosynthesis and extracellular release of tryptophan, enabling the pneumococcus to resist macrophage-mediated killing and reprogram the macrophage transcriptional landscape in a tryptophan-dependent manner. Disruption of tryptophan biosynthesis abolishes both colonization and invasive disease. Together, these findings reveal that mucin is not merely a barrier to be circumvented but an active signal that reprograms the pneumococcus toward a persistent, host-tolerant state. Our work herein identifies a distinctive repertoire of colonization mechanisms that allows a formidable pathogen to maintain long-term residence in its human host.

## RESULTS

### Mucin promotes prolonged pneumococcal stationary phase survival

The natural environment for the pneumococcus is the human nasopharynx which is replete with mucins. In the nasal passages, Muc5ac is the dominant mucin, followed by Muc5b, with low levels of Muc2. In addition to previously generated *Muc5ac* knockout mice^23^, we developed *Muc2* knockout mice (Fig. S1A, B) to ask the initial question if mucins impact pneumococcal colonization. Knockout and heterozygous mice were intranasally colonized with 10^5^ CFU of wild type BHN97 and the nasal bacterial population was monitored. *S. pneumoniae* BHN97 (serotype 19F) is a good model of nasal colonization and therefore used to investigate the impact of mucin on pneumococcal colonization^24^. While there were no differences in the initial colonization phenotypes at 24 hours post challenge, *Muc2*^−/−^ mice had a significant decrease in BHN97 nasal colonization at later time points (Fig.1A). In contrast, the *Muc5ac* knockout and heterozygous mice maintained a similar colonization by BHN97 throughout the period we monitored (Fig. 1B). These results provide evidence that not all mucins contribute similarly to pneumococcal biology; specifically, Muc2 plays an essential role in pneumococcal colonization *in vivo*.

**Figure 1.**
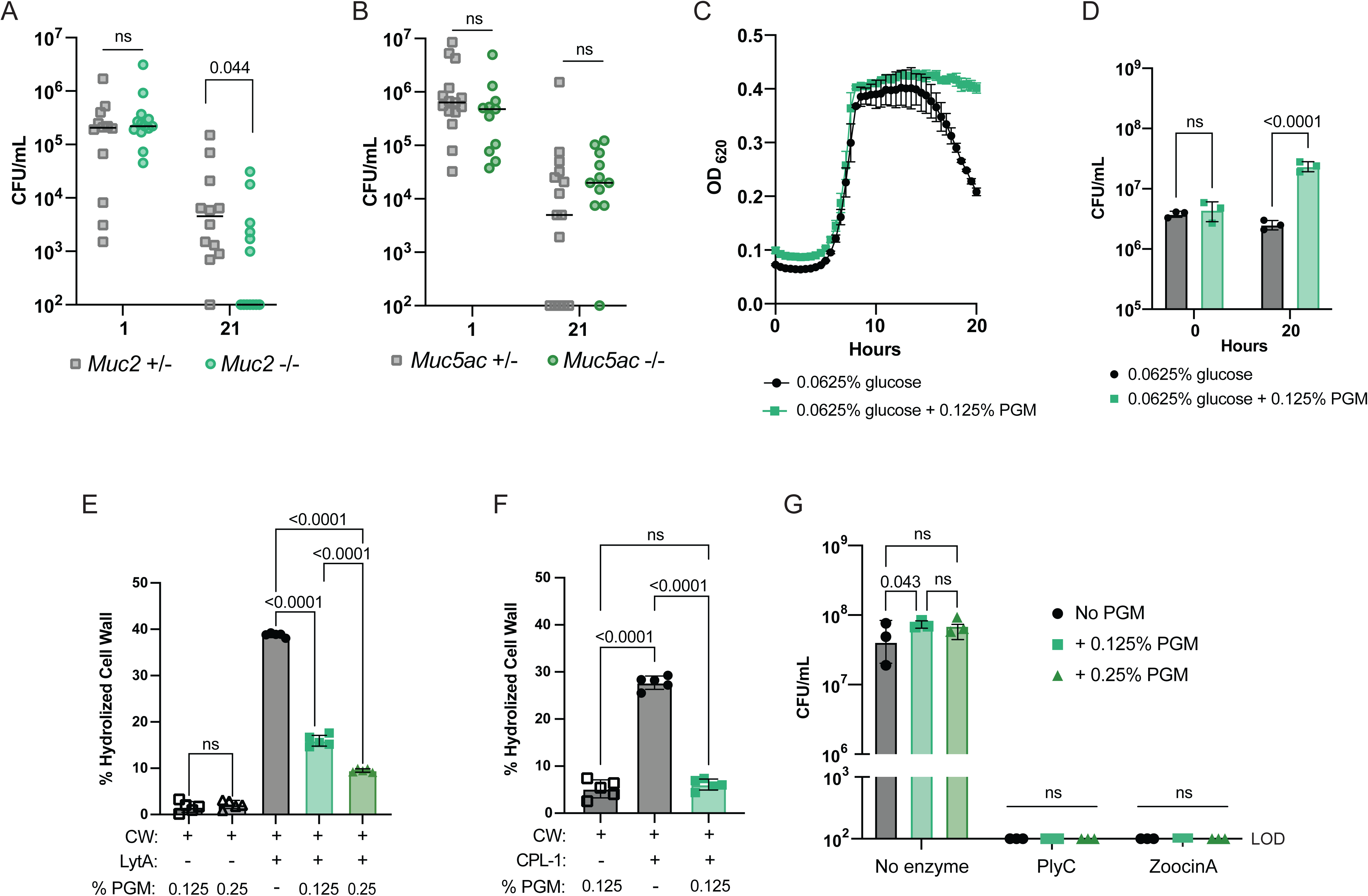
LytA-mediated autolysis is hindered when the pneumococcus is grown in media supplemented with porcine gastric mucin (PGM). *Muc2* +/− and *Muc2* −/− (A) and *Muc5ac* +/− and *Muc5ac* −/− (B) mice were intranasally colonized with 10^5^ CFU BHN97 and monitored. Bar represents median. Significance determined by Mann-Whitney test. (C) Media lacking carbon sources was supplemented with 0.0625% glucose ± 0.125% PGM and growth of *S. pneumoniae* BHN97 (serotype 19F) was followed for 20 hours. Mean and SD of biological duplicates. (D) At 20 hours, cells were plated for viability. Cell wall hydrolysis assay with purified rLytA (E) or CPL-1 (F) and cell wall, with or without PGM. Mean and SD of 5 biological replicates shown. Significance determined by one-way ANOVA. (G) *S. pyogenes* and *S. mutans* were incubated with recombinant PlyC or ZoocinA, respectively. Bacterial titers were determined 1 hourpost incubation. Mean and SD of biological triplicates graphed. Significance determined by 2-way ANOVA with Tukey’s multiple comparisons.

To determine the mechanisms behind how mucin impacts pneumococcal biology in the context of colonization, we first set out to determine if the model mucin porcine gastric mucin (PGM) could act as a carbon source for pneumococcal growth. Glucose and sucrose were replaced in the growth media and PGM was supplemented as the sole carbon source. There was no BHN97 growth with PGM as the sole carbon source (Fig. S2A). This shifted the focus to PGM supplementation in media containing glucose. We then followed the growth kinetics through stationary phase where the pneumococcus rapidly undergoes autolysis with a concomitant reduction in cellular viability^25^. Surprisingly, when pneumococcal strain BHN97 was grown in media containing 0.0625% glucose, PGM supplementation resulted in prolonged stationary phase survival of *S. pneumoniae* (Fig. 1C), with a survival benefit of several orders of magnitude following 20 hours of growth (Fig. 1D). This mucin-mediated survival was tested across multiple concentrations of PGM and found to be titratable (Fig. S2B). To determine if this was due to the osmotic stability and gel-like properties of the mucins preventing autolysis, we grew BHN97 in the presence of the fish gelatin and glycerol. Neither of these supplements prevented pneumococcal autolysis (Fig. S2C, D). Most pneumococci encode two neuraminidases^26^, NanA and NanB^27,28^. The pneumococcal neuraminidases are essential for colonization and have been demonstrated to interact with mucus^29–31^. To determine if the stationary phase survival was dependent on the interaction between the neuraminidases and sialic acid decorations on the mucin, we generated a Δ*nanAB* double knockout in BHN97 and grew it with 0.125% PGM. In wild type BHN97, we did not see a significant increase in expression of *nanA* or *nanB* in mucin conditions, and we recovered similar bacterial titers at 20 hours compared to wild type when grown with PGM (Fig. S2E, F). These data indicate that mucins promote stationary phase survival via a mechanism independent of osmotic protection or neuraminidase activity. As *S. pneumoniae* is characterized by a high degree of genetic variability as well as variation in the surface capsular polysaccharide, we sought to determine whether the inhibition of autolysis by mucin was a conserved feature across strains. We found the PGM-mediated inhibition of autolysis phenotype was observed across multiple pneumococcal strain backgrounds and capsule types (Fig. S3) suggesting the mucin mediated inhibition of autolysis is a conserved feature in *S. pneumoniae*.

### Mucin specifically interferes with the activity of pneumococcal cell wall amidases

Pneumococcal autolysis is driven by extracellular cleavage of peptidoglycan linkages via the cell wall amidase LytA. Early in stationary phase LytA performs its enzymatic activity, leading to cell autolysis, drop in optical density, and a rapid reduction in viable bacterial cells^32^. We hypothesized that the prolonged stationary phase and increased cell viability is due to loss of LytA enzymatic activity in the presence of PGM. To assess this, we took a biochemical approach using purified recombinant LytA and purified pneumococcal cell wall to quantify rLytA cell wall hydrolysis in the face of mucin^33^. In this assay, the enzymatic activity of LytA results in a rapid decrease in optical density as the pneumococcal cell wall is cleaved. When PGM was supplemented into the reaction, we observed significantly decreased cell wall hydrolysis by rLytA in a dose dependent manner (Fig. 1E). We next wanted to know if mucin interferes with the activity of other cell-wall degrading enzymes, or if it is specific to LytA. CPL-1 is a bacteriophage lysin that, like LytA, encodes a choline-binding domain that allows it to specifically bind to the pneumococcal cell wall and has been shown to lyse all tested pneumococcal strains^34,35^. We repeated the same assay performed with LytA and saw that CPL-1 is also inhibited when mucin is present (Fig. 1F). Next, we expanded our assay to assess the ability of mucin to interfere with cell wall degrading enzymes from other *Streptococcus* species. PlyC is a multimeric bacteriophage lysin that is active against group A streptococcal species^36,37^. Zoocin A is an endopeptidase encoded by *Streptococcus equi* that hydrolyzes the peptidoglycan bridge in susceptible species, including *Streptococcus mutans*^38,39^. When bacteria were challenged with their corresponding enzyme, we observed complete lysis of the culture, regardless of whether mucin was present (Fig. 1G). These data provide evidence that mucin specifically interferes with the activity of pneumococcal cell wall hydrolases, and not those of closely related upper airway colonizers.

### Pneumococci grown in mucins exhibit decreased cytotoxicity to human cells

Pneumolysin (PLY) is a cholesterol dependent cytolysin that is the predominant toxin produced by *S. pneumoniae*. Unique amongst the cholesterol dependent cytolysins, PLY lacks a signal sequence and largely relies upon autolysis for toxin release^40^. We hypothesized that if LytA lytic activity and subsequent autolysis were impaired, there should be a mucin-dependent decrease in the amount of released PLY and a corresponding decrease in cellular toxicity mediated by the pneumococcus. Supplementation of media with PGM resulted in a marked decrease in the amount of PLY released into the supernatant from strain BHN97 (Fig. S4A), while the loading control of all proteins recognized by human IgG is higher in the cells grown with mucin (Fig. S4B). As PLY release was compromised, we next sought to determine the impact of mucin supplementation on pneumococcal-mediated cytotoxicity. To test this, we incubated lung epithelial A549 cells with spent media of BHN97 grown in 0.25% PGM and measured cell viability via respiration measured by alamarBlue (Fig. S4C). Spent media of bacteria grown with PGM had significantly higher amount of A549 respiration, indicative of reduced cell death. Collectively these data indicate that mucin-mediated inhibition of autolysis restricts the release of PLY resulting in a corresponding decrease in host cell toxicity, validating previously published work with similar results from another pneumococcal strain^41^.

### Mucin is a signal for pneumococcal secretion of *de novo* synthesized tryptophan

The nasopharynx is a nutritionally limited environment^13,42^, and questions remain surrounding the metabolic constraints and adaptations the pneumococcus undergoes in this environment. Metabolomic profiling of the mouse nasopharynx during pneumococcal colonization has revealed that the upper airway metabolome is altered during carriage, with several amino acids elevated, though the origin of these changes was not determined^13^. Whether mucin, as the most abundant protein in mucus, could serve as a source of amino acids to support pneumococcal growth in this nutrient-limited niche is unknown. Therefore, we hypothesized that mucin could rescue pneumococcal growth in amino acid limited conditions. To this extent, we grew the pneumococcus in chemically defined media (CDM) lacking individual amino acids and compared it to growth in the same media supplemented with 0.125% PGM (Fig. S5, summarized in Fig. 2A). Interestingly, mucin rescued growth in all conditions where the pneumococcus exhibited a growth defect, except glutamine, irrespective of whether it encodes a biosynthetic operon for the amino acid or not. We were particularly interested in the amino acids that are known to be immunomodulatory, specifically arginine^42^, histidine^43^, leucine^44^, and tryptophan^45^. First, we wanted to determine if the pneumococcus was able to liberate these amino acids from mucin. We grew the pneumococcus in CDM lacking each amino acid, supplemented with mucin, for four hours. At this point we pelleted the cells, filtered the supernatant and measured the abundance of each amino acid via targeted mass spectrometry (Fig. 2B). For each amino acid, we observed an increase in free amino acid. Next, we wanted to investigate if the pneumococcus was able to synthesize the amino acids *de novo*. Here, we substituted the glucose in the CDM with ^13^C_6_ D-glucose to measure incorporation into *de novo* amino acids via mass spectrometry. Surprisingly, we found that *S. pneumoniae* grown in tryptophan deficient media supplemented with PGM produces a significantly higher amount extracellular labelled tryptophan as compared to conditions lacking mucin, suggesting this essential amino acid was being synthesized and released by the pneumococcus in response to mucin (Fig. 2C). Interestingly, the tryptophan biosynthetic locus, *trp*, was upregulated even when mucin was present in tryptophan rich media (Fig. 2D), suggesting that biosynthesis of tryptophan is not dependent on the lack of tryptophan in the environment. Therefore, mucin serves as a signal for the pneumococcus to upregulate tryptophan biosynthesis and release it into the environment, potentially impacting the host.

**Figure 2.**
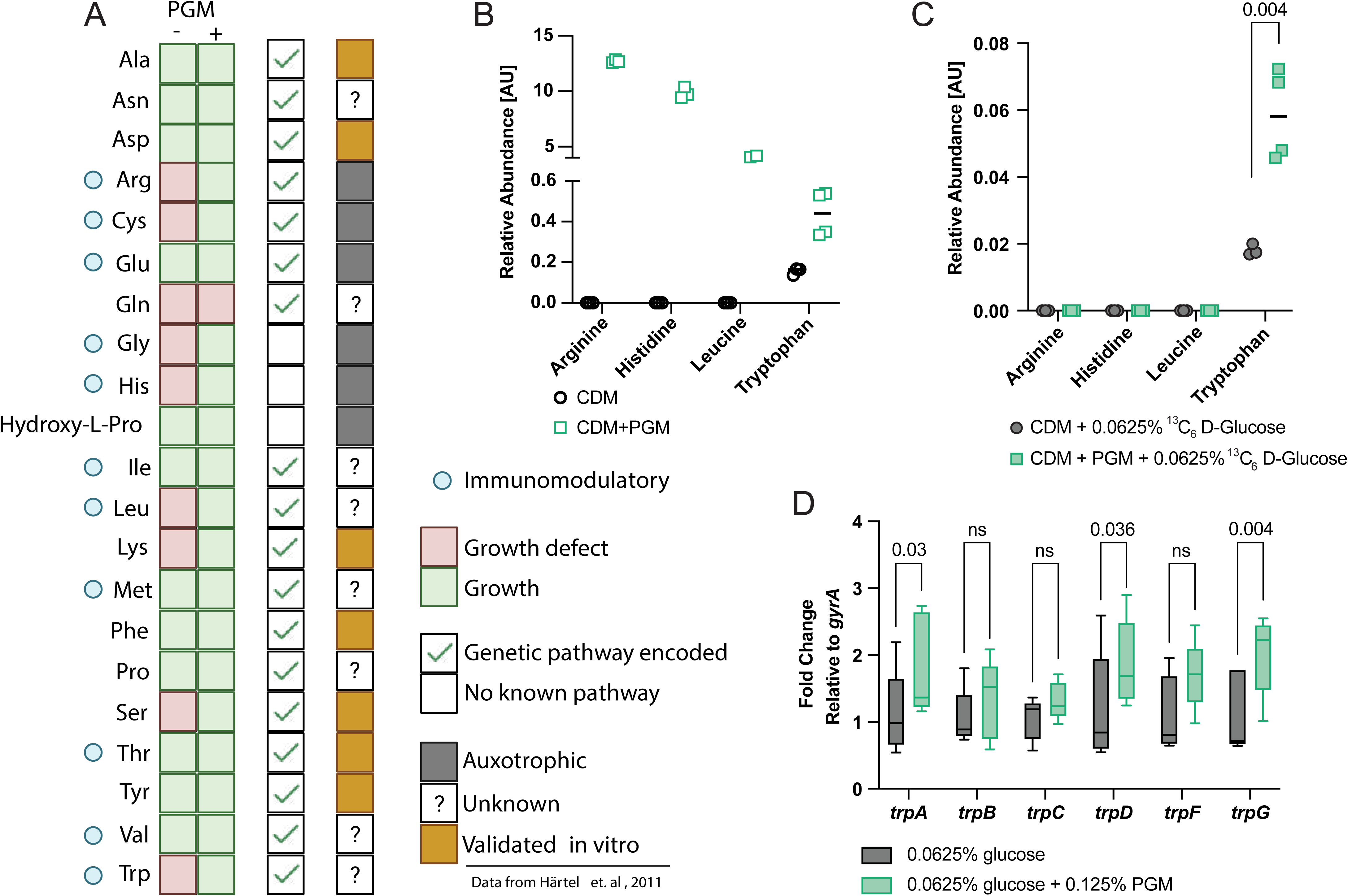
Mucus serves as an environmental cue to induce pneumococcal biosynthesis of tryptophan. (A) Mucin rescues pneumococcal growth defects in amino acid limited media. Summary of data shown in Fig. S4. (B) Quantification of selected amino acids by mass spectrometry in the supernatant of *S. pneumoniae* BHN97 grown with or without 0.125% PGM for 4 hours. (C) Quantification of heavy amino acids by mass spectrometry in the supernatant of *S. pneumoniae* BHN97 grown in media with ^13^C_6_ D-glucose as the sole carbon source, with or without 0.125% PGM for 4 hours. Significance determined by t test. (D) qPCR of the tryptone locus on BHN97 grown in complete media, including tryptophan, with or without 0.125% PGM. Box and whiskers plot of 5 replicates. Significance determined by 2-way ANOVA with Sidak’s multiple comparisons.

### Pneumococcal-derived tryptophan is necessary for survival against macrophages

The impact of tryptophan biosynthesis on pneumococcal colonization and invasive disease has not been defined, though several genetic screens have implicated a prominent role for this pathway at the host-pathogen interface^20,21,46^. To explore how tryptophan biosynthesis directly impacts pneumococcal colonization, we generated a knockout of the tryptophan biosynthesis locus, designated Δ*trp*, which was phenotypically a tryptophan auxotroph (Fig. S6A). This strain was used to colonize wild type C57BL/6 mice intranasally, and within three days was undetectable (Fig. 3A). We found that the Δ*trp* mutant grew comparably to wild type in mouse and human serum (Fig. S6B, C); however, the mutant demonstrated a growth defect in mouse and human whole blood (Fig. S6D, E). This led us to hypothesize that if the pneumococcus can synthesize tryptophan, then they are able to evade clearance by innate immune cells. Macrophages are the host’s first line of defense against pneumococcal insult^22,47^. Tryptophan can be metabolized into N-formylkynurenin via indoleamine 2,3-dioxygenase (IDO) and tryptophan 2,3-dioxygenase (TDO) in the kynurenine pathway. IDO expression in mononuclear phagocytes leads to expression of M2-specific markers, which are canonically associated with an anti-inflammatory macrophage phenotype^48^. Macrophages use tryptophan depletion as an antibacterial strategy, limiting the availability of the amino acid in the phagolysosome^49^. Therefore, we hypothesized that the *trp* locus is necessary for increased survival within macrophages. In an *in vitro* gentamycin protection assay with RAW 264.7 cells, pneumococci lacking the *trp* locus are recovered intracellularly from macrophages at a significantly lower rate than wild type bacteria at 30 minutes post gentamycin exposure (Fig. 3B). This phenotype is reversed in a the complemented BHN97 Ω*trp*. Unfortunately, BHN97 Ω*trp* overexpresses the *trp* locus and does not phenocopy wild type BHN97 in the *in vivo* colonization model (data not shown), likely due to a “goldilocks” level of expression that is beneficial for the pneumococcus. This is in accordance with a study whereby Warrier *et al*. recently reported that pneumococcal mutants with a constituently active regulator of the *trp* T-box complemented *in vitro* phenotypes, but not *in vivo*, demonstrating that too much expression is deleterious *in vivo*, too little is deleterious *in vivo* and *in vitro*, and wild type-level regulation is optimal^50^.

**Figure 3.**
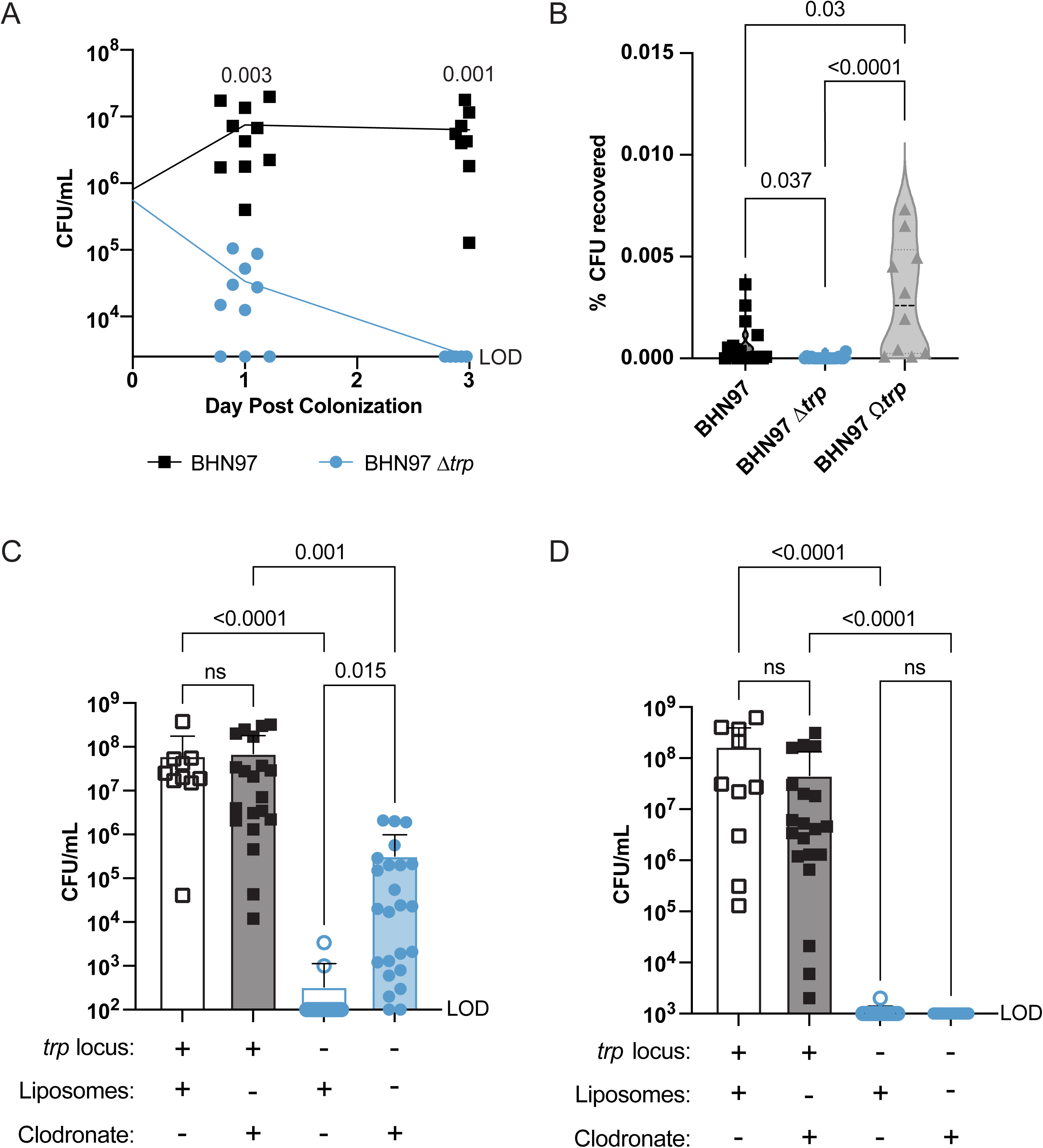
Tryptophan biosynthesis is essential for avoiding macrophage clearance in pneumococcal colonization and invasive disease. (A) Intranasal colonization of wild type BHN97 or an isogenic Δ*trp* biosynthesis operon knockout in wild type C57BL/6 mice. Significance determined by Mann-Whitney test at each time point. (B) RAW 264.7 cells were challenged with wild type, Δ*trp*, or a complemented Ω*trp* strain of BHN97 at an MOI of 100 for 30 minutes, at which point the media was replaced with media supplemented with gentamycin. 30 minutes after gentamycin exposure, macrophages were lysed and bacteria were enumerated and compared to input titers. Significance determined by Kruskal-Wallis test with Dunn’s multiple comparisons. C57BL/6 mice were treated with 100 µL clodronate or control liposomes intranasally 48 hours prior to infection with 10^6^ CFU wild type or a Δ*trp* pneumococcal mutant un 100 µL intranasally. 14 hours after infection, bacterial titers were enumerated from lungs (C) and blood (D). Significance determined by Kruskal-Wallis test with Dunn’s multiple comparisons. LOD = Limit of Detection.

While the pneumococcus is a commensal of the human nasopharynx, it has a proclivity for causing invasive infection. We next hypothesized that the *trp* operon is necessary for macrophage evasion *in vivo*. To investigate the impact of tryptophan biosynthesis on invasive pneumococcal infection, we generated a Δ*trp* mutant in TIGR4 (Fig. S6F). Like BHN97, the TIGR4 mutant does not exhibit a growth defect in mouse and human serum, but does in blood (Fig. S6G, H, I J). Additionally, the capacity of this mutant to cause invasive infection was severely attenuated with no detectable colonies in the blood (Fig.S7A), spleen (Fig. S7B), and kidneys (Fig. S7C). To explore the role of macrophages in the context of invasive disease, we administered clodronate into the lungs of wild type C57BL/6 mice to deplete the alveolar macrophages. 48 hours after clodronate, we administered 10^6^ CFU of wild type or Δ*trp* TIGR4 intranasally in 100 µL. Wild type pneumococci were recovered at high levels in clodronate and control liposome treated mice (Fig. 3C). Δ*trp* is detected only at low levels in liposome treated mice; however, if the mice are macrophage depleted, we recovered significantly more Δ*trp* mutant bacteria (Fig. 3C). Interestingly, in matched blood samples, we did not observe breakthrough of the Δ*trp* mutant into the bloodstream of clodronate treated mice, likely due to the presence of circulating and splenic macrophages in these mice (Fig. 3D). Collectively, we identify an important role of host mucins in modulation of pneumococcal autolytic capacity and metabolism, specifically tryptophan biosynthesis, that play critical roles for bacterial fitness during colonization and invasive disease.

### Rapid intracellular killing of a pneumococcal tryptophan auxotroph permits macrophage proliferative transcriptional reprogramming

We found that 30 minutes after the addition of gentamycin, the Δ*trp* mutant was cleared more rapidly than wild type, indicating that the inability to synthesize tryptophan compromises bacterial persistence in the macrophage intracellular niche (Fig. 3B). With this finding, we next sought to understand the response of the macrophages via transcriptomic profiling to identify metabolic changes that distinguish these two infection outcomes. We repeated the gentamycin protection assay but collected the macrophages for RNA extraction immediately after the addition of gentamycin, when we recover similar CFU for both genotypes (Fig. 4A). To characterize the transcriptional responses underlying the differential clearance phenotype, we performed bulk RNA-seq on uninfected macrophages, macrophages infected with wild type BHN97, and macrophages infected with the isogenic Δ*trp* mutant. Principal component analysis (PCA) of variance-stabilized expression values confirmed that biological replicates within each group clustered together and that the three conditions separated along the first two principal components (Fig. 4B). PC1 captured the primary axis of variation separating uninfected from infected samples, while PC2 distinguished wild type infected from mutant infected macrophages, indicating that the two infection conditions impose transcriptionally distinct programs on the macrophage.

**Figure 4.**
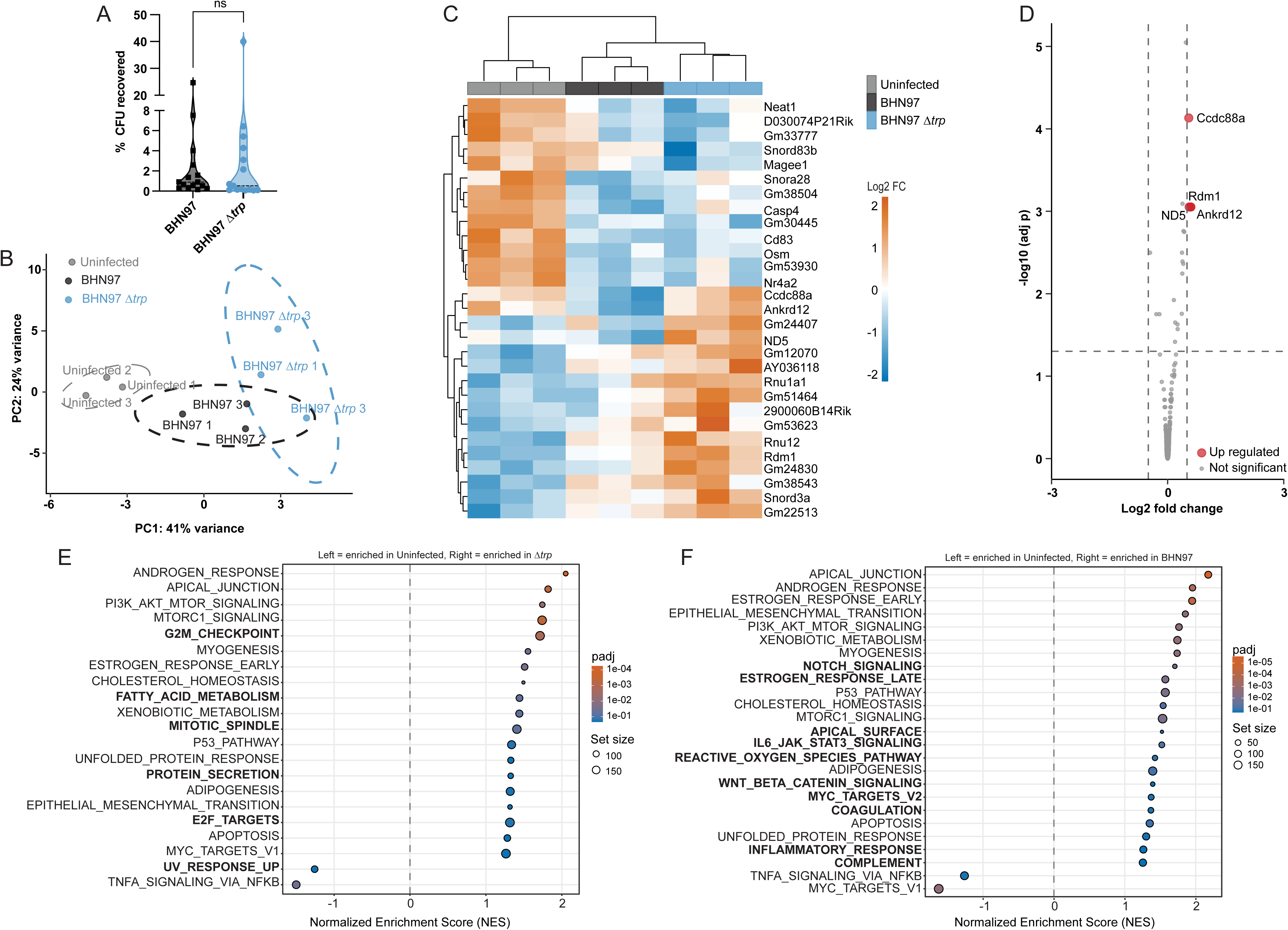
Tryptophan auxotrophy accelerates pneumococcal clearance and permits macrophage proliferative reprogramming. (A) RAW 264.7 cells were challenged with wild type or Δ*trp*, strain of BHN97 at an MOI of 100 for 30 minutes, macrophages were lysed and bacteria were enumerated and compared to input titers. Significance determined by Kruskal-Wallis test with Dunn’s multiple comparisons. (B) PCA plot of RNA sequencing data from uninfected macrophages or those infected with wild type BHN97 or Δ*trp* mutant. Ellipses represent 95% confidence. (C) Heatmap of the 29 significant differentially expressed genes. (D) Volcano plot of gene expression data reveals four genes upregulated in macrophages infected with Δ*trp* compared to wild type. GSEA pathway analysis using the Hallmark dataset reveals pathways that are upregulated in macrophages infected with Δ*trp* (E) or wild type (F) BHN97. Pathways unique to each condition are bolded.

Differential expression analysis (DESeq2, padj < 0.05, |log2FC| > 1.0) across the three conditions identified 29 differentially expressed genes in total (Fig. 4C). Unsupervised hierarchical clustering of these genes (z-scored VST values, Euclidean distance, complete linkage) revealed two distinct expression clusters. The cluster of 16 genes specifically induced in Δ*trp*-infected macrophages (uninfected mean z = −0.77, wild type mean z = −0.20, Δ*trp* mean z = +0.97), includes all four genes that reached individual significance in the volcano plot of genes differentially expressed between wild type and Δ*trp* (Fig. 4D: Ccdc88a, Rdm1, Ankrd12, ND5), as well as several small nuclear and nucleolar RNA genes (Rnu1a1, Rnu12, Snord3a) and predicted non-coding RNA loci. Ccdc88a (Girdin, log2FC = +0.55, padj = 7.4×10^−5^) encodes an actin-binding protein that physically interacts with NOD2 in macrophages to enhance bacterial sensing and clearance^51^. Ankrd12 (log2FC = +0.60, padj = 8.9×10^−4^) is a transcriptional corepressor that recruits histone deacetylases to nuclear receptor complexes^52^. Rdm1 (log2FC = +0.59, padj = 8.9×10^−4^) is an RNA recognition motif protein involved in DNA damage response^53^. ND5 (mt-Nd5, log2FC = +0.56, padj = 8.9×10^−4^) is a mitochondrially encoded Complex I subunit^54^. Complex I is a major source of mitochondrial ROS production, which contributes to macrophage bactericidal activity^55^. Per-group expression analysis revealed that for Ccdc88a and Ankrd12, wild type infection actively suppressed expression below the uninfected baseline (Ccdc88a: 740 (uninfected), 509 (wild type) counts; Ankrd12: 201 (uninfected), 115 (wild type) counts), while infection by the Δ*trp* mutant restored or slightly exceeded baseline levels (Ccdc88a: 794; Ankrd12: 210), suggesting that wild type BHN97 actively dampens these macrophage defense genes and the mutant cannot mount this suppression.

The modest quantity of individually significant genes in Fig. 4C, despite clear phenotypic divergence between the two infection conditions (Fig. 3), indicated that the transcriptional differences operate through coordinated, modest shifts across many genes rather than large-magnitude changes in a few, motivating us to explore pathway-level analysis by gene set enrichment analysis (GSEA). GSEA using the 50 pathways in the Hallmark collection was performed for each contrast using all ∼14,000 expressed genes ranked by Wald statistic, without applying a significance cutoff. This approach detects coordinated shifts across all genes in a pathway even when no individual gene reaches significance. Comparing the Δ*trp* vs uninfected (Fig. 4E) and wild type vs uninfected (Fig. 4F) GSEA results revealed that the two infection conditions share a core transcriptional response but diverge sharply on pathways governing proliferation, oxidative stress, and inflammatory signaling.

Several Hallmark pathways were enriched in both infection conditions relative to uninfected macrophages, representing a shared macrophage response to bacterial encounter. These included Apical Junction, Androgen Response, PI3K/AKT/mTOR Signaling, mTORC1 Signaling, Epithelial-Mesenchymal Transition, Xenobiotic Metabolism, Cholesterol Homeostasis, and Myogenesis. Both infections also suppressed TNFα Signaling via NFκB, indicating that dampening of acute inflammatory signaling is a shared feature of *S. pneumoniae* infection rather than a mutant-specific effect. The most striking differences between the two analyses emerged on pathways that shifted in opposite direction. MYC Targets V1 was induced by Δ*trp* infection but suppressed by wild type infection below the uninfected baseline. MYC is a master transcriptional factor that controls cell growth and proliferation^56^. Next, we looked at pathways that are unique to each infection (bolded). Pathways that were specifically activated in Δ*trp* infected macrophages include G2M Checkpoint, Fatty Acid Metabolism, Mitotic Spindle, Protein Secretion, and E2F Targets (Fig. 4E). Pathways specific for wild type infected macrophages include Notch Signaling, Estrogen Response Late, Apical Surface, IL6/Jak/Stat3 Signaling, ROS Pathway, Wnt/β-Catenin Pathway, Coagulation, Inflammatory Response, and Complement (Fig. 4F). The collective GSEA results support a model in which tryptophan biosynthesis determines whether macrophages adopt a proliferative/reparative phenotype (Δ*trp* infection) or a stressed and arrested bactericidal phenotype (wild type infection). The enrichment of MYC/E2F/G2M in the Δ*trp* dataset and ROS in the wild type dataset aligns with the known biology of macrophage proliferation during infection and the suppression of proliferation by proinflammatory signaling. Together, these results define a model in which wild type *S. pneumoniae* infection suppresses MYC-dependent transcriptional programs while inducing ROS and stress pathways, whereas the Δ*trp* mutant is unable to sustain intracellular persistence, rapidly cleared and permits macrophage engagement of proliferative transcriptional programs.

## DISCUSSION

Environmental context is critical to understanding the fundamental biology of pathogens for their capacity to both colonize and cause invasive disease. *Streptococcus pneumoniae* is a successful pathobiont due to its ability to asymptomatically colonize the human nasopharynx and disseminate into the lower respiratory tract causing community acquired pneumonia. The pneumococcus can be detected in 40–95% of infants and adults maintain carriage at low frequency for weeks at a time^57^. This prevalence argues that the pneumococcus has evolved not primarily to cause disease, but to persist within its host. Here we identify two mucin-mediated mechanisms that reframe the pneumococcus-human relationship as one of mutualism, in which both bacterium and host derive benefit from a stable colonization state (Fig. 5). Mucin, the defining protein of the nasopharyngeal environment, serves as the dual signal that suppresses autolysis and induces *de novo* tryptophan biosynthesis and release, positioning it as the molecular hinge between bacterial fitness and host tolerance.

**Figure 5.**
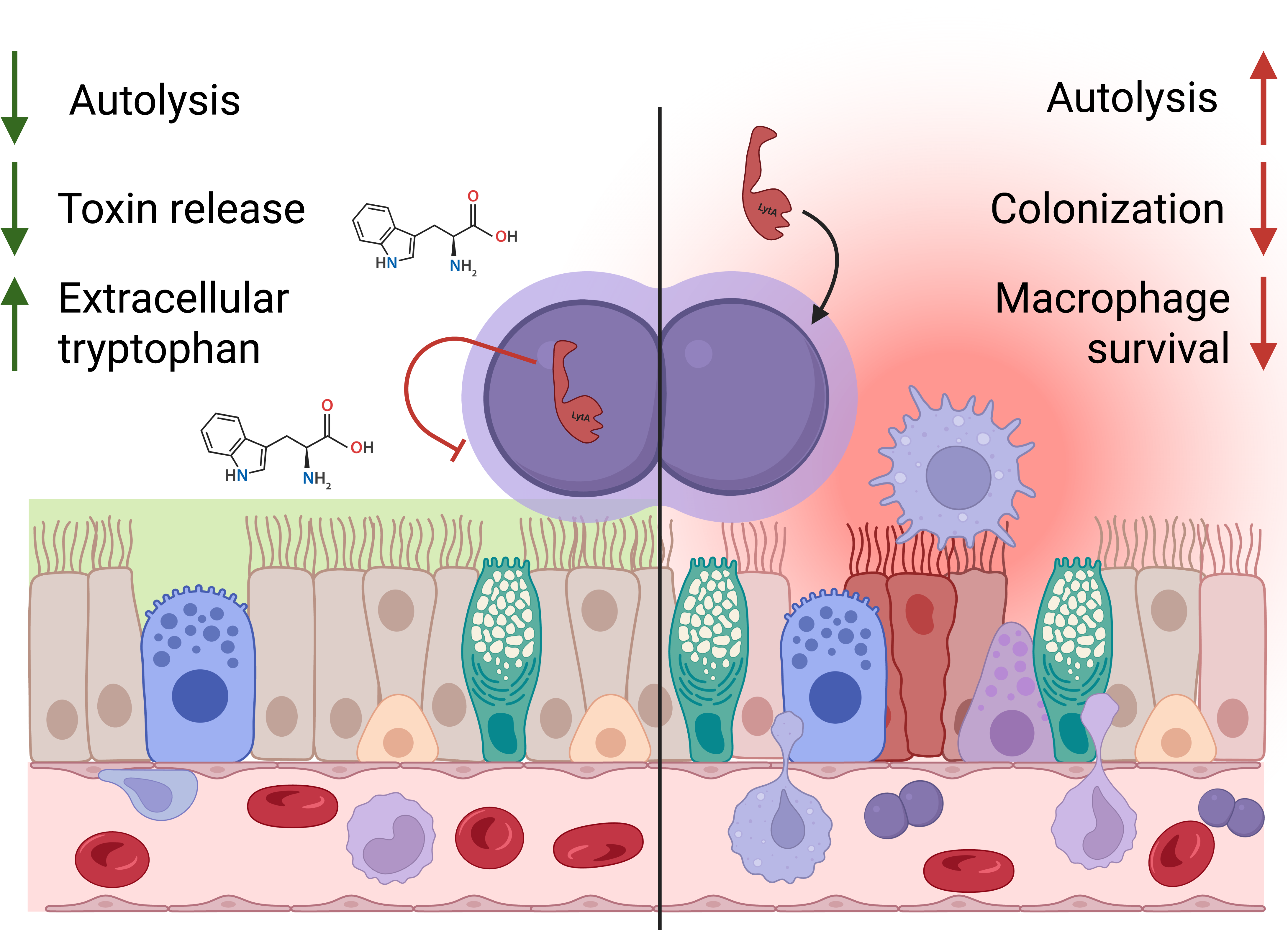
Mucin promotes pneumococcal colonization through decreased autolysis leading to decreased PLY release and promoting tryptophan synthesis and release, increasing survival against macrophages.

Our initial observations in mucin knockout mice led us to determine the mechanisms behind how mucins impact pneumococcal colonization (Fig. 1A, B). Autolysis is among the most studied features of pneumococcal biology. Driven by the amidase LytA, peptidoglycan linkages are cleaved and results in the characteristic optical density decline and viability loss in stationary phase^58^. LytA-mediated autolysis is the primary route by which pneumolysin (PLY), the pneumococcal cholesterol-dependent cytolysin, is released into the environment^59^. PLY is a potent toxin that integrates into host membranes, activates neutrophil reactive oxygen species production, and drives inflammation that characterizes invasive pneumococcal disease. Our data show that mucin specifically inhibits the enzymatic activity of LytA and the pneumococcal phage lysin CPL-1 while leaving unrelated cell wall hydrolases unaffected (Fig. 1C-G). This mucin specificity indicates that mucin does not act as a generic physical barrier to enzymatic activity but interferes with a particular class of pneumococcal cell wall hydrolases, providing a targeted survival benefit. In this light, mucin-mediated autolysis suppression is not simply a bacterial persistence mechanism, it is a mutualistic switch. The bacterium gains extended viability in a nutrient-limited niche while the host is spared the cytolytic and inflammatory consequences of PLY exposure (Fig. S4). Notably, sub-lytic concentrations of PLY have been shown to trigger host repair and anti-inflammatory responses rather than damage^60^, suggesting that the residual PLY released under mucin-suppressed conditions may actively contribute to a tolerogenic environment rather than a destructive one.

The second mucin-responsive mechanism we identified extends the mutualistic logic from bacterial survival to active immune modulation. The nasopharynx is a nutritionally limited environment, and the ability of the pneumococcus to liberate amino acids from mucin is consistent with the metabolic adaptations required for sustained carriage (Fig. 2A, B). What was unexpected was the finding that mucin acts not merely as a substrate but as a signal. The *trp* biosynthetic locus is upregulated in the presence of mucin even in tryptophan-replete media (Fig. 2D), and the pneumococcus synthesizes and releases significantly more tryptophan when mucin is present (Fig 2C). Tryptophan is unique among the amino acids in its centrality to host-microbe mutualism. It is an essential amino acid in humans and must be obtained from diet or bacteria that possess the tryptophan biosynthetic pathway^61^. Once acquired, tryptophan can be metabolized into a diverse array of bioactive derivatives. In humans, approximately 85% of available tryptophan is catabolized through the kynurenine pathway via IDO and TDO, ∼5% enters the serotonin pathway, and the remainder is converted by microbiota into indole and other derivatives^62^. Kynurenine pathway metabolites drive macrophage polarization toward an anti-inflammatory M2 phenotype and promote immune tolerance^63^, while indole derivatives engage the aryl hydrocarbon receptor, a critical interface between microbiota and host mucosal immunity that balances regulatory T cell and Th17 responses and promotes barrier integrity and tissue repair^18,64^. In the gut, microbial tryptophan metabolism through these pathways is now recognized as a cornerstone of mucosal homeostasis^65–68^. The pneumococcus, by synthesizing and releasing tryptophan in response to mucin, is provisioning the host with an essential metabolite, in a niche where dietary tryptophan may be scarce. The host is then able to convert tryptophan into derivatives that calibrate the local immune response toward tolerance. This is mutualism; the bacterium supplies a metabolite the host needs but cannot make, and the host metabolism of that metabolite creates an environment that permits bacterial persistence.

Our macrophage infection and transcriptomic data provide functional support for this model (Fig. 3, 4). Macrophages use tryptophan depletion as an antibacterial strategy, catabolizing the amino acid via IDO to starve auxotrophic organisms in the phagolysosome^15,69^. The pneumococcus, by retaining its own biosynthetic capacity, can resist this nutritional immunity. Consistent with this, the Δ*trp* mutant is rapidly cleared from macrophages and is undetectable *in vivo* within three days of colonization, demonstrating that tryptophan biosynthesis is essential for intracellular persistence and colonization fitness (Fig. 3. RNAseq of infected macrophages revealed that the two infection conditions impose transcriptionally distinct programs (Fig. 4). At the individual gene level, wild type BHN97 actively suppresses macrophage defense genes below uninfected baseline (Fig. 4B). Ccdc88a (Girdin), which enhances NOD2-mediated bacterial sensing and clearance, and Ankrd12, a transcriptional corepressor in nuclear receptor pathways, were both reduced by wild type infection below uninfected levels, while the Δ*trp* mutant failed to mount this suppression. GSEA revealed the broader architecture of these divergent responses. The Δ*trp* mutant permitted engagement of proliferative and reparative transcriptional programs (MYC targets, E2F targets, G2M checkpoint, mitotic spindle), consistent with a macrophage that has successfully cleared the bacterium and transitioned to a reparative phase (Fig. 4E). Wild type infection specifically activated IL6/Jak/Stat3 signaling, Inflammatory Response, Complement, and ROS pathways, illustrating a bactericidal, stress-arrested macrophage state (Fig. 4F). Together, these data paint a picture in which tryptophan biosynthesis does not simply suppress macrophage defenses but enables the pneumococcus to survive a bactericidal macrophage state while preventing the macrophage from completing its clearance program and transitioning to proliferative tissue repair.

This mutualistic relationship aligns with a growing appreciation that the nasopharyngeal microbiome is not merely a reservoir of potential pathogens but a structured community whose members co-evolve with the host to support mucosal immunity^70,71^. The host benefit we identify operates at two levels. Mucin-mediated autolysis suppression directly reduces pneumococcal cytotoxicity to human epithelial cells, and tryptophan release provides the host with an essential amino acid that serves as substrate for immunomodulatory derivative production. The macrophage transcriptomic data further show that wild type pneumococci prevent the macrophage from completing clearance and transitioning to proliferative tissue repair, permitting bacterial persistence without triggering the destructive remodeling associated with reparative proliferation. In both mechanisms, the pneumococcus limits the very activities that would damage the host and trigger clearance, while providing something the host needs.

Several limitations of this study should be acknowledged. First, our *in vitro* experiments rely on porcine gastric mucin (PGM), which differs in glycosylation and protein composition from the human airway mucins that the pneumococcus encounters in its natural niche. Second, the macrophage transcriptional data were generated in an *in vitro* RAW 264.7 infection model, which cannot fully recapitulate the cellular diversity, spatial organization, and mucosal context of the nasopharyngeal immune environment *in vivo*. The macrophage proliferative and bactericidal programs we identified should be validated in primary airway macrophages and in the context of intact mucosal tissue. Finally, while our data demonstrate that the pneumococcus synthesizes and releases tryptophan in response to mucin and that tryptophan biosynthesis is required for macrophage evasion, the direct links between pneumococcal-derived tryptophan and host IDO/TDO activity, kynurenine production, M2 polarization, and AhR-mediated tolerance remain inferential.

The pneumococcus has spent far more of its evolutionary history as a nasopharyngeal colonizer than as an invasive pathogen, and this is reflected in its biology. In the presence of mucin, the pneumococcus curtails its own cytotoxicity while provisioning the host with an essential amino acid that the host cannot synthesize and that feeds into immunomodulatory pathways promoting mucosal tolerance. Mucin serves as the environmental signal that coordinates this mutualistic program, linking bacterial metabolism to host immune modulation. Viewing the pneumococcus through this lens does not diminish the seriousness of pneumococcal disease but rather begins to reframe infection as a failure of colonization homeostasis.

## MATERIALS AND METHODS

### Bacterial strains, growth conditions, and generating mutants

Pneumococcal strains were grown on tryptic soy agar (EMD Chemicals) supplemented with 3% defibrinated sheep blood and 20μg/mL neomycin. In all experiments expect for cells grown in chemically defined media (CDM, see below), pneumococcal cultures were inoculated from newly streaked TSA blood plates into complete C+Y media^72^ lacking sucrose, supplemented with 0.0625% glucose, and grown at 37°C, 5% CO_2_. When indicated, media was supplemented varying amounts (w/v) of porcine gastric mucin (PGM, Sigma). Growth curves were performed in a 96 well plate in a Biotek Synergy, with starting OD_620_ of ∼0.05 for all strains. For growth in human serum (Sigma), mouse serum (Abcam), human blood (Gulf Coast Blood), or mouse blood pneumococci were grown to an OD_620_ of 0.1 in C+Y, and 10uL was added to 1mL serum or blood, grown under standard conditions, and enumerated at various timepoints.

Allelic replacements were made by replacing the gene of interest with an antibiotic resistance cassette by splicing overlap extension PCR. A region 1 kB upstream and downstream of the gene of interest was amplified by PCR with Takara HotStart polymerase according to manufacturer instructions from parental genomic DNA using the primers listed below with an overhang corresponding to the beginning and/or end of the antibiotic resistance cassette. The upstream and downstream fragments were mixed with the antibiotic cassette and amplified with Takara polymerase and upstream forward and downstream reverse primer. The resulting PCR product was gel purified using the MinElute kit (Qiagen) according to manufacturer’s instructions. BHN97 was transformed with the purified PCR product in C+Y media using both CSP-1 and CSP-2, while TIGR4 was transformed with CSP-2. The tryptophan biosynthesis operon was knocked out of *S. pneumoniae* strains BHN97 (serotype 19) and TIGR4 (serotype 4) using the same primers and method. Briefly, genes *trpA* (SP_1811), *trpB* (SP_1812), *trpF* (SP_1813), *trpC* (SP_1814), *trpD* (SP_1815), *trpG* (SP_1816), and *trpE* (SP_1817) were replaced with an erythromycin resistance cassette. Deletions were confirmed with colony PCR of the *trp* locus. To generate the *nanAnanB* double mutant in BHN97, *nanA* was interrupted with an erythromycin resistance cassette and *nanB* was interrupted with a kanamycin resistance cassette and confirmed via colony PCR.

### C+Y media

Media was prepared as previously published^24^:

Supplement: Prepare the supplement “3 in 1” salts by adding 50 g MgCl_2_ 6H_2_O, 0.25 g CalCl_2_ anhydrous and 0.1 ml (0.1 M) Manganese sulfate 4 H2O in 500 ml distilled water (dH_2_O), mix well and autoclave. Prepare 500 mL of 20% Glucose, 50% Sucrose, 2mg/ml Adenosine and 2mg/ml Uridine 500 each respectively, and filter to sterilize, store at 4°C. Combine all 5 components at the following ratio: 60 ml “3 in 1” salts, 120 mL 20% Glucose, 6 mL 50% Sucrose, 120ml 2mg/mL Adenosine, 120 mL 2mg/mL Uridine, mix in a beaker and filter sterilize, label as Supplement, store at 4°C.

Prepare Adams I by combining the following chemicals: 30 mg Nicotinic Acid (Niacin stored at 40C), 35 mg Pyridoxine HCl (B6), 120 mg Ca-Pantothenate (stored at 4°C), 32 mg Thiamine-HCl, 14 mg Riboflavin, and 0.06 mL Biotin (0.5mg/mL stock). Add dH_2_O to 200mL, then add 1–5 drops of 10N NaOH to dissolve chemicals, filter sterilized and store in foiled bottles at 4°C.

Prepare Adams II by adding the following chemicals: 50 mg FeSO4•7H2O, 50 mg CuSO_4_, 50 mg ZnSO4•7H2O, 20 mg MnCl_2_ and 1ml HCl (concentrate), up to 100ml dH_2_O, filter to sterilize, store at 4°C.

Prepare Adams III by adding the following 5 components: 800 mg Asparagine, 80 mg Choline Chloride, 64 mL Adams I, 16 mL Adams II and 0.64 ml CaCl_2_ (1% stock), to 400 ml dH_2_O. Filter sterilize solutions and store in foiled bottle at 4°C.

Buffers: Prepare 1M KH_2_PO_4_ and 1M K_2_HPO_4_ (autoclaved) as the stocks, mix 26.5 mL 1M KH_2_PO_4_ and 473 mL 1M K_2_HPO_4_ and stir well, do not titrate, filter to sterilize, 4°C.

PreC: Mix 4.83 g Sodium Acetate (Anhydrous), 20 g Difco Casamino Acids/technical, 20 mg/L-Tryptophan, and 200 g/L Cysteine HCl, dissolve in 800 dH_2_O, adjust pH to 7.4–7.6 by adding 10 N NaOH, stir well for 60 minutes, fill up to 4 liter dH_2_O, mix well, aliquot 400 mL portions in 500 mL flasks, autoclave for 30 minutes, and store at 4°C.

C+Y: Add 0.5g glutamine to 500 mL dH_2_O, filter to sterilize and store at 4°C. Add 2 g pyruvic acid (stored at 4°C) to 100 ml dH_2_O, filter to sterilize, and store at 4°C. Solve 5g yeast to 100mL dH_2_O (25g in 500mL), and autoclave (filter to sterilize if necessary). Add the following solutions to 400 mL PreC: 13 mlLSupplement, 10 mL Glutamine, 10 mL Adams III, 5 mL Pyruvate, 15 mL K-Phosphate buffer, and 9 mL Yeast. Filter sterilize and store at 4°C. If preparing C+Y that lacks glucose and sucrose, substitute with an equal volume of dH_2_O.

### Primers

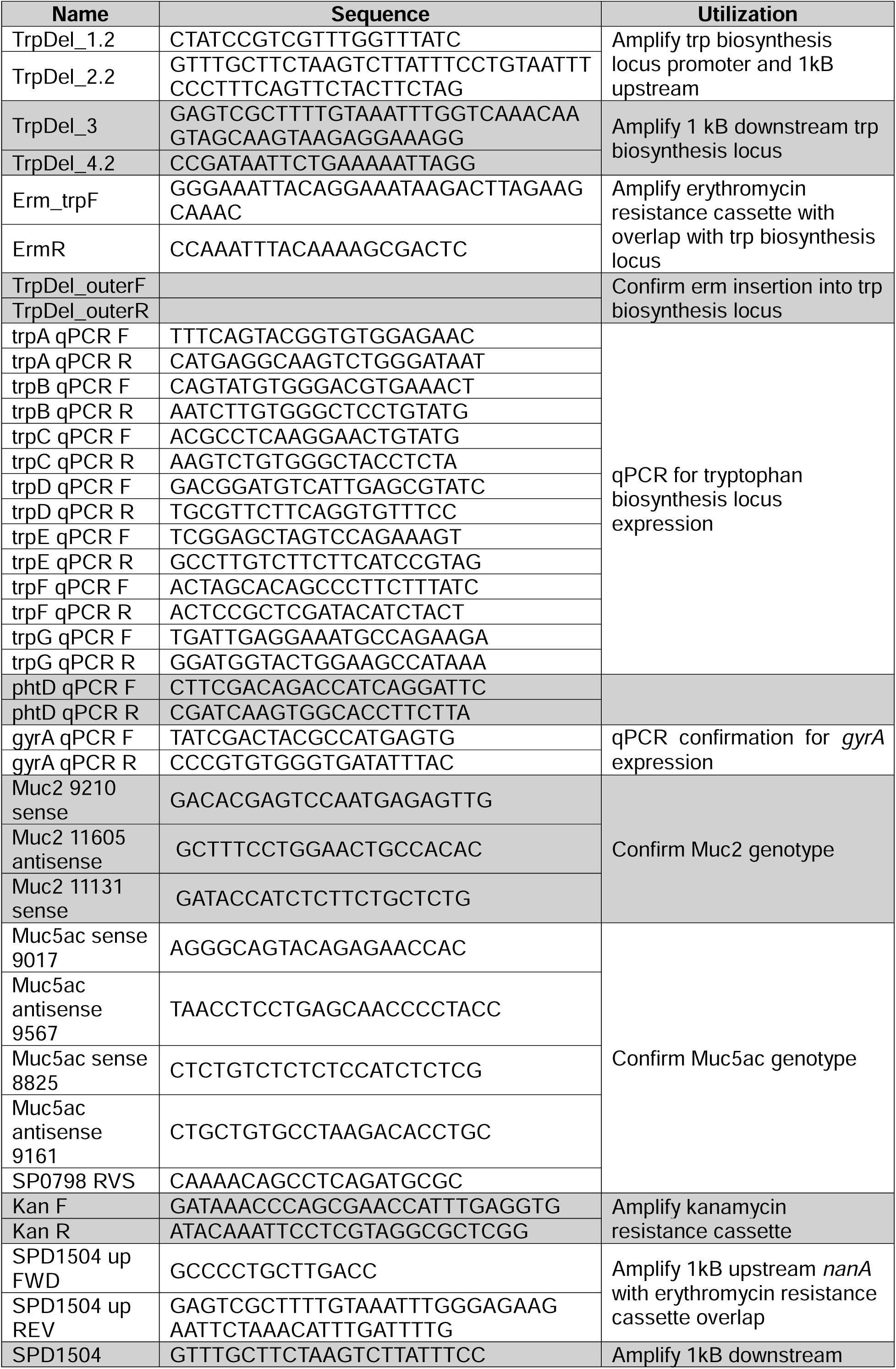
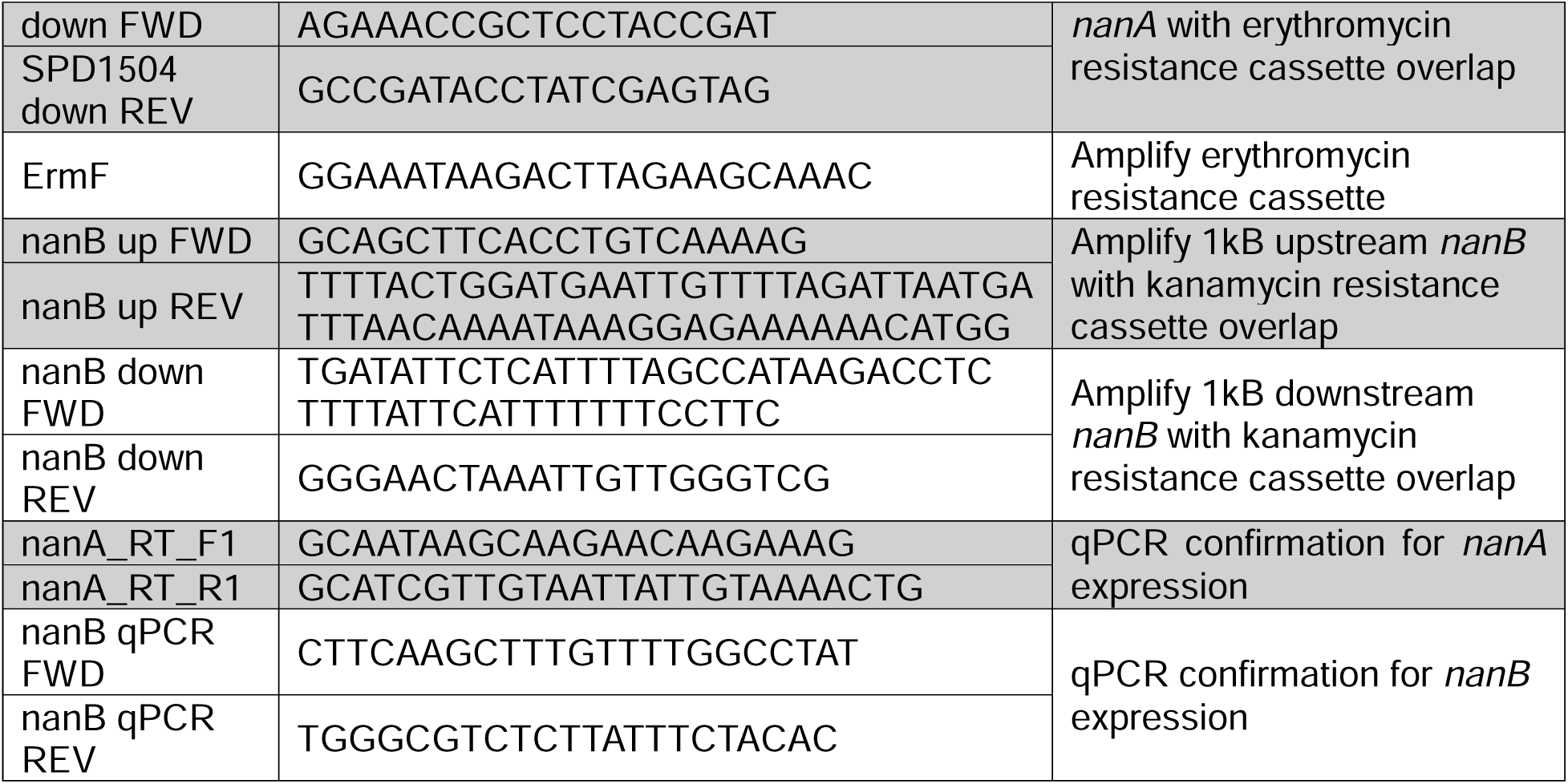

### Cell wall hydrolysis assay

Recombinant pneumococcal LytA or CPL-1 was expressed in and purified from *E. coli* using constructs and methods as previously described^33,73^. Cell wall was purified from *S. pneumoniae* R6 following previously described methods^74^. Cell wall was diluted 1:100 in PBS and sonicated for 30 minutes. 28nM rLytA or 100nM CPL-1 was added to 1mL cell wall, with or without 0.125% or 0.25% PGM resuspended overnight in PBS. The reaction was incubated at 37°C, 5% CO_2_ for 15 minutes. OD_620_ was measured and the percent hydrolyzed cell wall was calculated by comparing to control cell wall conditions that lacked rLytA or CPL-1 at the same timepoint. Five biological replicates were performed.

### Lysis of streptococcal species with purified enzymes

*Streptococcus pygoenes* HSC5 and *Streptococcus mutans* UA159 were grown in Todd-Hewitt broth (Difco) supplemented with 0.2% yeast extract (Gibco) to an OD_620_ of 1, then back diluted to 0.6. 1 mL of cells were pelleted and resuspended in 20 mM Tris + 3 mM CaCl_2_. 100 µL *S. pyogenes* were incubated with 2 µL PlyC (4.5 mg/mL), or 100 µL *S. mutans* were incubated with 2 µL ZoocinA (4 mg/mL). PGM was dissolved in the Tris+CaCl_2_ buffer and added to achieve 0.125% or 0.25%. Bacteria were incubated at 37°C for 1 hour, after which cells were enumerated via plating. Purified recombinant PlyC and ZoocinA were graciously provided by the Korotkova Lab at the University of Kentucky.

### Western blots for pneumococcal proteins

*S. pneumoniae* TIGR4 (serotype 4) or BHN97 (serotype 19F) was grown under designated conditions for 6 hours at which point 1 mL was taken from the culture and the cells were pelleted. Supernatant was saved and cells were washed in PBS once. Cell pellets were lysed with 10% SDS and 10% DOC, following previously published protocols^75^. Lysed cells and protein precipitates boiled for 15 minutes in 1x loading dye (NuPAGE, Invitrogen). 15 µL of sample were added to each well of a 10 well 4-12% Bis-Tris precast protein gel (Invitrogen) and ran at 100V for 80 minutes. Proteins were transferred to nitrocellulose membranes and blocked in 5% non-fat milk/PBS-Tween20. Rabbit polyclonal anti-PLY^75^ or 10% Human IgG (Grifols, NDC 13533-800-20) primary antibodies were diluted to 1:3000 in 5% non-fat milk PBS-Tween20 and incubated at room temperature for 1 hour or 4°C overnight. Membranes were washed 3 times in 0.1% PBS-Tween20. Appropriate secondary HRP-conjugated antibody (Bio-Rad) was incubated at 1:10,000 in 5% non-fat milk/PBS-Tween20 at room temperature for 1 hour, followed by 3 washes in 0.1% PBS-Tween20. Membranes were developed with Dura (Thermo Scientific) for 5 minutes and imaged on the ChemiDoc imager (Bio-Rad).

### A549 cell respiration quantification

Cytotoxicity of PGM grown pneumococcal cells on A549 cells was performed following a previously published protocol by Bath *et al*^76^. A549 cells (ATCC CCL-185) were grown under standard conditions at 37°C with 5% CO_2_. *S. pneumoniae* TIGR4 was resuspended to an OD_620_ of 0.2 and grown for 3 hours in denoted media. Bacterial cells were pelleted and spent media was passed through 0.22 µM filters to remove bacterial cells. Spent media, along with fresh media controls, were added to A549 cells that were suspended at 10^5^ cells per well in a 96-well plate in technical triplicate. alamarBlue HS cell viability reagent (Invitrogen) was added to the plates and incubated at 37°C with 5% CO_2_ for 1 hour. Fluorescence was measured at 560/590 nm using Biotek. Percent cell respiration was calculated relative to fresh media control.

### Generation of Muc2 −/− mice

A *Muc2* knockout mouse (*Muc2*^−/−^) was generated using CRISPR technology to delete exon 1 of *Muc2* in C57BL/6J in the Regional Mouse Genetics Core facility at UC Denver. Guide RNA (gRNA) design was done using a combination of CRISPOR and the Broad Institute sgRNA^77,78^. Guide RNA targets 9470rev (5’-ACCTCACTTCACGTCATCCA in the 5’ flanking region, PAM located 534 bp upstream of the *Muc2* start codon in Exon 1) and 11269rev (5’-CTATCCCACATGCCCCCCA in intron 2, PAM located 229 bp downstream of Exon 2) were chosen as to disrupt *Muc2*.

Injection was performed in zygotes generated by IVF using C57BL/6J sperm and C57BL/6N eggs. Zygotes were then transferred into pseudopregnant recipients. F0 pups were genotyped by PCR using primers outside the region to be modified to identify putative positive founders. Screening was performed using Muc2 9210 sense and Muc2 11605 antisense primers located outside the gRNA target regions, which allowed for amplification of wild-type *Muc2* (2,396 bp) and disrupted *Muc2* (593 bp). Sequencing confirmed deletion and sites of non-homologous end-joining.

Mice were subsequently bred using a heterozygous breeding strategy with C56BL6/J for outcrossing >5 generations. For routine genotyping assays, *Muc2* 9210 sense and *Muc2* 11605 antisense were used for detecting knockout alleles as above. For wild type detection and *Muc2* 11131 sense was used in combination with *Muc2* 11605 antisense. *Muc2* absence was verified in small and large intestine tissue samples by immunohistochemistry.

### Pneumococcal intranasal murine colonization and infection

For all mouse experiments, male and female mice were used. Genotype was confirmed using primers listed in the above table. Knockout and heterozygous control mice are littermates. For experiments, mice were 8-12 weeks old, and age was randomized between genotypes and sex and experimental groups.

For pneumococcal colonization, 10^5^ pneumococcal cells were administered intranasally to isoflurane anesthetized mice in 25 µL sterile PBS. No prior antibiotic treatment was administered to the mice. To monitor bacterial colonization, mice were anesthetized with isoflurane and 25 µL sterile PBS was administered to the mouse nostrils and immediately removed. Pneumococci were enumerated via selective plating on TSA blood agar plates supplemented with 20 µg/mL neomycin, as is standard for pneumococcal selection. For intraperitoneal (IP) infection, 100 CFU (lethal dose killing 100%)^79^ of wild type TIGR4 or an isogenic tryptophan biosynthesis mutant were administered to mice IP in 100 µL PBS. At 21 hours post infection, mice were euthanized, and blood, kidneys, and spleen were collected. Organs were homogenized in sterile PBS and pneumococci were enumerated by selective plating on blood agar plates with 20 µg/mL neomycin.

All experiments involving animals were performed with the prior approval of and in accordance with the guidelines of the St. Jude Institutional Animal Care and Use Committee (Protocol 538, approved 10 June 2022). The St. Jude laboratory animal facilities are fully accredited by the American Association for Accreditation of Laboratory Animal Care. The laboratory animals are maintained in accordance with the applicable portions of the Animal Welfare Act and the guidelines prescribed in the DHHS publication Guide for the Care and Use of Laboratory Animals.

### Clodronate depletion of alveolar macrophages

Clodrosomes or control empty liposomes (Encapsula Nano Sciences) were diluted 1:1 in PBS and 100 µL was administered intranasally to anesthetized mice following a previously published protocol^80^. 48 hours after depletion, mice were infected intranasally with 10^6^ CFU of *S. pneumoniae* TIGR4 or an isogenic mutant lacking the tryptophan biosynthesis pathway diluted in 100 µL of PBS. 24 hours after infection, mice were euthanized and pneumococci were enumerated from lungs via selective plating.

### Chemically defined media

| Chemical | Solvent | Concentration (mg/L) |
|---|---|---|
| **100x Salt A** |  |  |
| K<sub>2</sub>HPO<sub>4</sub> | Water | 200 |
| KH<sub>2</sub>PO<sub>4</sub> | Water | 500 |
| **50x Salt B** |  |  |
| MgSO<sub>4</sub>•7H<sub>2</sub>O | Water | 700 |
| FeSO<sub>4</sub>•7H<sub>2</sub>O | Water | 5.04 |
| Fe(NO<sub>3</sub>)<sub>3</sub>•9H<sub>2</sub>O | Water | 1.06 |
| MnSO<sub>4</sub>•H<sub>2</sub>O | Water | 5.58 |
| **100x Amino Acids – individually prepared** |  |  |
| L-Alanine | Water | 100 |
| L-Arginine | Water | 100 |
| L-Aspartic Acid | 0.1M NaOH | 100 |
| L-Asparagine | Water | 100 |
| L-Cysteine | 0.1M NaOH | 500 |
| L-Cystine | 0.2M HCl | 50 |
| L-Glutamic Acid | 0.1M NaOH | 100 |
| Glycine | Water | 100 |
| L-Histidine | Water | 100 |
| L-Isoleucine | Water | 100 |
| L-Leucine | Water | 100 |
| L-Lysine | Water | 100 |
| L-Methionine | Water | 100 |
| L-Phenylalanine | Water | 100 |
| L-Proline | Water | 100 |
| Hydroxy-L-Proline (trans) | Water | 100 |
| L-Serine | Water | 100 |
| L-Threonine | Water | 100 |
| L-Tryptophan | 0.5M HCl | 100 |
| L-Tyrosine | 0.1M NaOH | 100 |
| L-Valine | Water | 100 |
| **100x Vitamins Group A** |  |  |
| p-Aminobenzoic Acid | Water | 0.2 |
| Biotin | Water | 0.2 |
| Pyridoxamine dihydrochloride | Water | 1 |
| **100x Vitamins Group B** |  |  |
| Niacinamide | Water | 1 |
| B-Nicotinamide adeninedinucleotide | Water | 2.5 |
| Pantothenate calcium salt | Water | 2 |
| **100x Vitamins Group C** |  |  |
| Pyridoxal HCl | Water | 1 |
| Thiamine hydrochloride | Water | 1 |
| Vitamin B12 | Water | 0.1 |
| **200x Folic Acid** |  |  |
| Folic Acid | 0.1M NaOH | 0.8 |
| **100x Riboflavin** |  |  |
| Riboflavin | 0.1M NaOH | 1.6 |
| **50x Nucleotides** |  |  |
| Adenine | 0.2M NaOH | 20 |
| Guanine | 0.2M NaOH | 20 |
| Uracil | 0.2M NaOH | 20 |
| **20x Individual Components** |  |  |
| CaCl<sub>2</sub>•2H<sub>2</sub>O | Water | 20 |
| NaC<sub>2</sub>H<sub>3</sub>O<sub>2</sub>•3H<sub>2</sub>O | Water | 2,720 |
| NaHCO<sub>3</sub> | Water | 2,500 |
| NaH<sub>2</sub>PO<sub>4</sub>•H<sub>2</sub>O | Water | 3,200 |
| C<sub>3</sub>H<sub>14</sub>NOCl | Water | 20 |
| Na<sub>2</sub>HPO<sub>4</sub>•2H<sub>2</sub>O | N/A | 9,220 |

For 100 mL media, add 38.5 mL water, and each of the above components to achieve a 1x concentration. Media should have a final pH of 7.4. Glucose can be added to a final concentration of 0.0625% when necessary. Filter sterilize media, store at room temperature and use within 3 days.

### Heavy glucose tracing to measure tryptophan production

BHN97 was grown in chemically defined media (CDM) lacking glucose and tryptophan. ^13^C_6_ D-glucose (Cambridge Isotope Laboratories) was provided as a carbon source. When indicated, 0.125% PGM was added to the media. Pneumococci were grown for 4 hours at which point 1 mL culture was pelleted and cells were enumerated. Media was removed and stored at –20°C until analysis was performed.

Methanol was added to media to a final concentration of 80% and incubated at –80°C for 1 hr. Samples were centrifuged at 3500 x g for 10 min to remove cell debris. The supernatant was removed to a new glass tube and was dried overnight using a Savant SP1010 SpeedVac. Samples were resuspended in 80% methanol and analyzed using a Shimadzu Prominence Ultra-Fast Liquid Chromatograph (UFLC) attached to a Sciex QTrap 4500 system equipped with a Turbo V ion source (Sciex). Samples were injected into an XSelect HSS C18 column (2.5 µm pore size, 3.0 by 150 mm) using a flow rate of 0.3 ml/min. Solvent A contained 100 mM ammonium formate (pH 5.0), 2% acetonitrile, and 0.1% tributylamine. Solvent B was composed of 95% acetonitrile, 50 mM ammonium formate (pH 6.3), and 0.1% tribuylamine. The HPLC program was as follows: starting solvent mixture of 0% solvent B, 0 to 2 min isocratic with 0% solvent B; 2 to 12 min linear gradient to 5% solvent B; 12 to 17 min linear gradient to 90% solvent B; 17 to 25 min isocratic with 90% solvent B; 25 to 27 min linear gradient to 0% solvent B; 27 to 30 min isocratic with 0% solvent B. The Sciex QTrap 4500 system was operated in the positive mode. The ion source parameters were as follows: ion spray voltage, 5,500 V; curtain gas pressure, 20 psi; temperature, 400°C; collision gas setting, high; ion source gas 1 pressure, 25 psi; ion source gas 2 pressure, 40 psi; declustering potential, 10 V; and collision energy, 18 V. The multiple-reaction monitoring (MRM) for tryptophan and its stable isotope forms are: Trp, 205/188; +2-Trp, 207/190; +4-Trp, 209/192; +6-Trp, 211/194; +8-Trp, 213/196; +9-Trp, 214/197; and +11-Trp, 216/199. The system was controlled by the Analyst software (Sciex) and analyzed with Multi-Quant 3.0.2 software (Sciex).

### Macrophage gentamycin protection assay

RAW-Blue macrophage cells (Invitrogen), derived from RAW 264.7 macrophage cells, were maintained following ATCC recommendation in DMEM (Gibco) supplemented with 10% sterile FBS (Corning) under the standard conditions of 37°C with 5% CO_2_. For the gentamycin protection assay, cells were seeded at 5×10^4^ cells per well in a 24 well plate (Corning). The next day, pneumococci were grown in C+Y to an OD of 0.4, pelleted and resuspended in sterile PBS, then diluted to a concentration that is equivalent to an MOI of 100 in DMEM+FBS, calculated by averaging the total number of macrophages per well from 3 wells. Input pneumococci were enumerated by plating. Macrophages were incubated with the pneumococci for 30 mins, at which point media was removed and replaced with DMEM+FBS supplemented with 200ug/mL gentamycin (Gibco). After 30 mins, media was removed, macrophages were lysed in cold 0.025% triton-x (Sigma), and pneumococci were enumerated.

### Macrophage RNAseq

The macrophage gentamycin protection assay was repeated, and immediately after gentamycin addition, macrophages were collected in Trizol (Invitrogen). To obtain sufficient RNA, 5 wells were combined for each biological replicate. RNA was extracted via bead beating and following the Qiagen RNeasy kit. Libraries were prepared for sequencing following the NEB Ultra II RNA Library Kit for Illumina with the Unique Dual Inde Adaptors and Primers. Libraries were sequenced by the Hartwell Center at St. Jude Children’s Research Hospital.

Bulk RNA-seq reads were aligned to the mouse GRCm39 reference genome (Ensembl release 110 annotation) using HISAT2 with reverse-stranded library configuration^81^. Gene-level read counts were quantified with featureCounts^82^ (strand-specific mode, –s 2). Differential expression analysis was performed in R using DESeq2^83^ with apeglm^84^ log2 fold-change shrinkage, across three comparisons: WT vs Neg (uninfected), Δ*trp* vs Neg, and Δ*trp* vs WT. Genes meeting adjusted p < 0.05 and |log2FC| > 1.0 were deemed significant. For visualization, variance-stabilizing transformation (VST) was applied, and significant genes were hierarchically clustered using pheatmap (Euclidean distance, complete linkage) with optimal cluster number determined by silhouette width. Gene set enrichment analysis was conducted with fgsea^85^ against the MSigDB Hallmark gene set collection (50 pathways; *Mus musculus* orthologs accessed via msigdbr)^86^, with genes ranked by Wald statistic (log2FC / lfcSE) and significance defined as adjusted p < 0.25. Enrichment dotplots and running-score plots were generated with ggplot2^87^. Computations were performed on the St. Jude Children’s Research Hospital High-Performance Computing Facility and analysis was assisted by Biomni (Phylo)^88^. Sequence reads and gene expression data can be accessed from NCBI under PRJNA1525725 and GSE346916, respectively.

### Statistical analyses

All statistical analyses were performed in GraphPad Prism version 10.3.1. In each figure legend we report the statistical method that was performed.

## ACKNOWLEDGEMENTS

C.N.J. is supported by NIH grants T32-AI106700 and F32-AI286447. C.M.E. and C.A. are supported by NIH R03 DK114545. We would like to thank the Genetically Engineered Murine Models Core at the University of Colorado Anschutz School of Medicine for design and generation of the *Muc2*^−/−^ mouse strain. This research included experiments conducted by the Hartwell Center for Bioinformatics & Biotechnology, which is supported, in part, by ALSAC and National Cancer Institute grant P30 CA021765.

## REFERENCES

1 Robb, C. T., Regan, K. H., Dorward, D. A. & Rossi, A. G. Key mechanisms governing resolution of lung inflammation. Semin Immunopathol 38, 425–448 (2016). 10.1007/s00281-016-0560-6

2 Pneumococcus: Vaccine preventable diseases surveillance standards. (Publications of the World Health Organization, 2018).

3 Collaborators, G. B. D. L. R. I. Estimates of the global, regional, and national morbidity, mortality, and aetiologies of lower respiratory infections in 195 countries, 1990-2016: a systematic analysis for the Global Burden of Disease Study 2016. The Lancet Infectious Diseases 18, 1191–1210 (2018). 10.1016/S1473-3099(18)30310-4

4 Goldblatt, D. et al. Antibody responses to nasopharyngeal carriage of *Streptococcus pneumoniae* in adults: a longitudinal household study. J Infect Dis 192, 387–393 (2005). 10.1086/431524

5 Simell, B. et al. The fundamental link between pneumococcal carriage and disease. Expert Rev Vaccines 11, 841–855 (2012). 10.1586/erv.12.53

6 Witten, J., Samad, T. & Ribbeck, K. Selective permeability of mucus barriers. Curr Opin Biotechnol 52, 124–133 (2018). 10.1016/j.copbio.2018.03.010

7 McShane, A. et al. Mucus. Curr Biol 31, R938–R945 (2021). 10.1016/j.cub.2021.06.093

8 Ualiyeva, S. et al. A nasal cell atlas reveals heterogeneity of tuft cells and their role in directing olfactory stem cell proliferation. Sci Immunol 9, eabq4341 (2024). 10.1126/sciimmunol.abq4341

9 Nelson, A. L. et al. Capsule enhances pneumococcal colonization by limiting mucus-mediated clearance. Infect Immun 75, 83–90 (2007). 10.1128/iai.01475-06

10 Li, Y., Weinberger, D. M., Thompson, C. M., Trzciński, K. & Lipsitch, M. Surface charge of *Streptococcus pneumoniae* predicts serotype distribution. Infection and Immunity 81, 4519–4524 (2013). doi:10.1128/iai.00724-13

11 Glover, J. S., Ticer, T. D. & Engevik, M. A. Characterizing the mucin-degrading capacity of the human gut microbiota. Scientific Reports 12, 8456 (2022). 10.1038/s41598-022-11819-z

12 Tsuboi, Y. et al. Pattern changes of mucin gene expression with pneumococcal otitis media. International Journal of Pediatric Otorhinolaryngology 61, 23–30 (2001). 10.1016/S0165-5876(01)00540-7

13 Green, A. E. et al. Airway metabolic profiling during *Streptococcus pneumoniae* infection identifies branched chain amino acids as signatures of upper airway colonisation. PLOS Pathogens 19, e1011630 (2023). 10.1371/journal.ppat.1011630

14 Niño-Castro, A. et al. The IDO1-induced kynurenines play a major role in the antimicrobial effect of human myeloid cells against *Listeria monocytogenes*. Innate Immun 20, 401–411 (2014). 10.1177/1753425913496442

15 Schmidt, S. V. & Schultze, J. L. New insights into IDO biology in bacterial and viral infections. Front Immunol 5, 384 (2014). 10.3389/fimmu.2014.00384

16 Wang, X.-F. et al. The role of indoleamine 2,3-dioxygenase (IDO) in immune tolerance: focus on macrophage polarization of THP-1 cells. Cellular Immunology 289, 42–48 (2014). 10.1016/j.cellimm.2014.02.005

17 Zelante, T. et al. Tryptophan catabolites from microbiota engage aryl hydrocarbon receptor and balance mucosal reactivity via interleukin-22. Immunity 39, 372–385 (2013). 10.1016/j.immuni.2013.08.003

18 Stockinger, B., Shah, K. & Wincent, E. AHR in the intestinal microenvironment: safeguarding barrier function. Nat Rev Gastroenterol Hepatol 18, 559–570 (2021). 10.1038/s41575-021-00430-8

19 Li, Y. et al. Tryptophan and the innate intestinal immunity: crosstalk between metabolites, host innate immune cells, and microbiota. European Journal of Immunology 52, 856–868 (2022). 10.1002/eji.202149401

20 Jensen, P. A., Zhu, Z. & van Opijnen, T. Antibiotics disrupt coordination between transcriptional and phenotypic stress responses in pathogenic bacteria. Cell Reports 20, 1705–1716 (2017). 10.1016/j.celrep.2017.07.062

21 Torres, M., Paszti, S. & Eberl, L. Shedding light on bacteria–host interactions with the aid of TnSeq approaches. mBio 15, e00390–00324 (2024). doi:10.1128/mbio.00390-24

22 Lemon, J. K., Miller, M. R. & Weiser, J. N. Sensing of interleukin-1 cytokines during *Streptococcus pneumoniae* colonization contributes to macrophage recruitment and bacterial clearance. Infection and immunity 83, 3204–3212 (2015).

23 Hasnain, S. Z. et al. Muc5ac: a critical component mediating the rejection of enteric nematodes. Journal of Experimental Medicine 208, 893–900 (2011). 10.1084/jem.20102057

24 Rowe, H. M. et al. Bacterial factors required for transmission of *Streptococcus pneumoniae* in mammalian hosts. Cell Host Microbe 25, 884–891.e886 (2019). 10.1016/j.chom.2019.04.012

25 Mosser, J. & Tomasz, A. Choline-containing teichoic acid as a structural component of pneumococcal cell wall and its role in sensitivity to lysis by an autolytic enzyme. Journal of Biological Chemistry 245, 287–298 (1970).

26 Pettigrew, M. M., Fennie, K. P., York, M. P., Daniels, J. & Ghaffar, F. Variation in the presence of neuraminidase genes among *Streptococcus pneumoniae* isolates with identical sequence types. Infection and immunity 74, 3360–3365 (2006).

27 Gut, H., King, S. J. & Walsh, M. A. Structural and functional studies of *Streptococcus pneumoniae* neuraminidase B: an intramolecular trans-sialidase. FEBS letters 582, 3348–3352 (2008).

28 McCombs, J. E. & Kohler, J. J. Pneumococcal neuraminidase substrates identified through comparative proteomics enabled by chemoselective labeling. Bioconjugate chemistry 27, 1013–1022 (2016).

29 Montgomery Matthew, T., Ortigoza, M., Loomis, C. & Weiser Jeffrey, N. Neuraminidase-mediated enhancement of *Streptococcus pneumoniae* colonization is associated with altered mucus characteristics and distribution. mBio 16, e02579–02524 (2024). 10.1128/mbio.02579-24

30 Marion, C., Burnaugh Amanda, M., Woodiga Shireen, A. & King Samantha, J. Sialic acid transport contributes to pneumococcal colonization. Infection and Immunity 79, 1262–1269 (2011). 10.1128/iai.00832-10

31 King, S. Pneumococcal modification of host sugars: a major contributor to colonization of the human airway? Molecular oral microbiology 25, 15–24 (2010).

32 García, E. Structure, function, and regulation of LytA: the N-acetylmuramoyl-l-alanine amidase driving the “suicidal tendencies” of *Streptococcus pneumoniae*—A review. Microorganisms 13, 827 (2025).

33 Kietzman, C. C., Gao, G., Mann, B., Myers, L. & Tuomanen, E. I. Dynamic capsule restructuring by the main pneumococcal autolysin LytA in response to the epithelium. Nat Commun 7, 10859 (2016). 10.1038/ncomms10859

34 Djurkovic, S., Loeffler, J. M. & Fischetti, V. A. Synergistic killing of *Streptococcus pneumoniae* with the bacteriophage lytic enzyme Cpl-1 and penicillin or gentamicin depends on the level of penicillin resistance. Antimicrobial Agents and Chemotherapy 49, 1225–1228 (2005). doi:10.1128/aac.49.3.1225-1228.2005

35 Grandgirard, D., Loeffler, J. M., Fischetti, V. A. & Leib, S. L. Phage lytic enzyme Cpl-1 for antibacterial therapy in experimental pneumococcal meningitis. The Journal of Infectious Diseases 197, 1519–1522 (2008). 10.1086/587942

36 Nelson, D., Schuch, R., Chahales, P., Zhu, S. & Fischetti, V. A. PlyC: a multimeric bacteriophage lysin. Proceedings of the National Academy of Sciences 103, 10765–10770 (2006).

37 King, H. et al. Molecular basis for recognition of the Group A Carbohydrate backbone by the PlyC streptococcal bacteriophage endolysin. Biochemical Journal 478, 2385–2397 (2021).

38 Gargis, S. R. et al. Zif, the zoocin A immunity factor, is a FemABX-like immunity protein with a novel mode of action. Applied and environmental microbiology 75, 6205–6210 (2009).

39 Rahman, M. M. et al. Glycosylation of serine/threonine-rich intrinsically disordered regions of membrane-associated proteins in streptococci. Nature Communications 16, 4011 (2025). 10.1038/s41467-025-58692-8

40 Martner, A., Dahlgren, C., Paton, J. C. & Wold, A. E. Pneumolysin released during *Streptococcus pneumoniae* autolysis is a potent activator of intracellular oxygen radical production in neutrophils. Infect Immun 76, 4079–4087 (2008). 10.1128/iai.01747-07

41 Bath, J. et al. Mucins protect against *Streptococcus pneumoniae* virulence by suppressing pneumolysin expression. The Journal of Clinical Investigation 134 (2024).

42 Krismer, B. et al. Nutrient limitation governs *Staphylococcus aureus* metabolism and niche adaptation in the human nose. PLOS Pathogens 10, e1003862 (2014). 10.1371/journal.ppat.1003862

43 Bäumer, W. & Roßbach, K. Histamine as an immunomodulator. JDDG: Journal der Deutschen Dermatologischen Gesellschaft 8, 495–504 (2010).

44 Bonvini, A., Coqueiro, A. Y., Tirapegui, J., Calder, P. C. & Rogero, M. M. Immunomodulatory role of branched-chain amino acids. Nutrition reviews 76, 840–856 (2018).

45 Moffett, J. R. & Namboodiri, M. A. Tryptophan and the immune response. Immunology and cell biology 81, 247–265 (2003).

46 van Opijnen, T. & Camilli, A. A fine scale phenotype-genotype virulence map of a bacterial pathogen. Genome Res 22, 2541–2551 (2012). 10.1101/gr.137430.112

47 Dockrell, D. H. et al. Alveolar macrophage apoptosis contributes to pneumococcal clearance in a resolving model of pulmonary infection. The Journal of Immunology 171, 5380–5388 (2003). 10.4049/jimmunol.171.10.5380

48 Sheu, K. M. & Hoffmann, A. Functional hallmarks of healthy macrophage responses: their regulatory basis and disease relevance. Annual Review of Immunology 40, 295–321 (2022). 10.1146/annurev-immunol-101320-031555

49 Sweet, M. J., Ramnath, D., Singhal, A. & Kapetanovic, R. Inducible antibacterial responses in macrophages. Nature Reviews Immunology 25, 92–107 (2025).

50 Warrier, I. et al. RNA cis-regulators are important for *Streptococcus pneumoniae in vivo* success. PLOS Genetics 20, e1011188 (2024). 10.1371/journal.pgen.1011188

51 Katkar, G. D. et al. Distinct colitis-associated macrophages drive NOD2-dependent bacterial sensing and gut homeostasis. The Journal of Clinical Investigation 135 (2025). 10.1172/JCI190851

52 Bai, R. et al. Clinical significance of Ankyrin repeat domain 12 expression in colorectal cancer. J Exp Clin Cancer Res 32, 35 (2013). 10.1186/1756-9966-32-35

53 Hamimes, S. et al. RDM1, a novel RNA recognition motif (RRM)-containing protein involved in the cell response to cisplatin in vertebrates. J Biol Chem 280, 9225–9235 (2005). 10.1074/jbc.M412874200

54 Bourges, I. et al. Structural organization of mitochondrial human complex I: role of the ND4 and ND5 mitochondria-encoded subunits and interaction with prohibitin. Biochemical Journal 383, 491–499 (2004).

55 West, A. P. et al. TLR signalling augments macrophage bactericidal activity through mitochondrial ROS. Nature 472, 476–480 (2011). 10.1038/nature09973

56 Dang, C. V. et al. in Seminars in cancer biology. 253–264 (Elsevier).

57 Weiser, J. N., Ferreira, D. M. & Paton, J. C. *Streptococcus pneumoniae*: transmission, colonization and invasion. Nature Reviews Microbiology 16, 355–367 (2018). 10.1038/s41579-018-0001-8

58 García, E. Structure, function, and regulation of LytA: the N-acetylmuramoyl-l-alanine amidase driving the “suicidal tendencies” of *Streptococcus pneumoniae*-a review. Microorganisms 13 (2025). 10.3390/microorganisms13040827

59 Nishimoto, A. T., Rosch, J. W. & Tuomanen, E. I. Pneumolysin: pathogenesis and therapeutic target. Frontiers in Microbiology Volume 11 – 2020 (2020). 10.3389/fmicb.2020.01543

60 Pereira, J. M., Xu, S., Leong, J. M. & Sousa, S. The yin and yang of pneumolysin during pneumococcal infection. Frontiers in Immunology Volume 13 – 2022 (2022). 10.3389/fimmu.2022.878244

61 McCann, J. R. & Rawls, J. F. Essential amino acid metabolites as chemical mediators of host-microbe interaction in the gut. Annu Rev Microbiol 77, 479–497 (2023). 10.1146/annurev-micro-032421-111819

62 Holeček, M. Serotonin, kynurenine, and indole pathways of tryptophan metabolism in humans in health and disease. Nutrients 18, 507 (2026).

63 Wang, X. F. et al. The role of indoleamine 2,3-dioxygenase (IDO) in immune tolerance: focus on macrophage polarization of THP-1 cells. Cell Immunol 289, 42–48 (2014). 10.1016/j.cellimm.2014.02.005

64 Zelante, T. et al. Tryptophan catabolites from microbiota engage aryl hydrocarbon receptor and balance mucosal reactivity via interleukin-22. Immunity 39, 372–385 (2013). 10.1016/j.immuni.2013.08.003

65 Gao, J. et al. Impact of the gut microbiota on intestinal immunity mediated by tryptophan metabolism. Frontiers in Cellular and Infection Microbiology Volume 8 – 2018 (2018). 10.3389/fcimb.2018.00013

66 Han, H., Safe, S., Jayaraman, A. & Chapkin, R. S. Diet-host-microbiota interactions shape aryl hydrocarbon receptor ligand production to modulate intestinal homeostasis. Annu Rev Nutr 41, 455–478 (2021). 10.1146/annurev-nutr-043020-090050

67 Li, Y. et al. Tryptophan and the innate intestinal immunity: crosstalk between metabolites, host innate immune cells, and microbiota. Eur J Immunol 52, 856–868 (2022). 10.1002/eji.202149401

68 Su, X., Gao, Y. & Yang, R. Gut microbiota-derived tryptophan metabolites maintain gut and systemic homeostasis. Cells 11 (2022). 10.3390/cells11152296

69 Niño-Castro, A. et al. The IDO1-induced kynurenines play a major role in the antimicrobial effect of human myeloid cells against *Listeria* i. Innate Immun 20, 401–411 (2014). 10.1177/1753425913496442

70 Yu, K. et al. Interactions between bacteria in the human nasopharynx: a scoping review. Lancet Microbe 6, 101062 (2025). 10.1016/j.lanmic.2024.101062

71 Cuthbertson, L. et al. Genomic attributes of airway commensal bacteria and mucosa. Commun Biol 7, 171 (2024). 10.1038/s42003-024-05840-3

72 Lacks, S. & Hotchkiss, R. D. A study of the genetic material determining an enzyme activity in pneumococcus. Biochimica et biophysica acta 39, 508–518 (1960).

73 Sanz, J. M., Lopez, R. & Garcia, J. L. Structural requirements of choline derivatives for ‘conversion’of pneumococcal amidase A new single-step procedure for purification of this autolysin. FEBS letters 232, 308–312 (1988).

74 Mann, B., Loh, L. N., Gao, G. & Tuomanen, E. Preparation of purified Gram-positive bacterial cell wall and detection in placenta and fetal tissues. Bio Protoc 6 (2016). 10.21769/BioProtoc.2037

75 Echlin, H., Frank, M., Rock, C. & Rosch, J. W. Role of the pyruvate metabolic network on carbohydrate metabolism and virulence in *Streptococcus pneumoniae*. Molecular Microbiology 114, 536–552 (2020). 10.1111/mmi.14557

76 Bath, J. et al. Mucins protect against *Streptococcus pneumoniae* virulence by suppressing pneumolysin expression. J Clin Invest 134 (2024). 10.1172/JCI182769

77 Doench, J. G. et al. Optimized sgRNA design to maximize activity and minimize off-target effects of CRISPR-Cas9. Nat Biotechnol 34, 184–191 (2016). 10.1038/nbt.3437

78 Concordet, J. P. & Haeussler, M. CRISPOR: intuitive guide selection for CRISPR/Cas9 genome editing experiments and screens. Nucleic Acids Res 46, W242–w245 (2018). 10.1093/nar/gky354

79 Gianfaldoni, C. et al. Sortase A confers protection against *Streptococcus pneumoniae* in mice. Infection and Immunity 77, 2957–2961 (2009). doi:10.1128/iai.01516-08

80 Nishimoto, A. T. et al. RNA quality control enables antibiotic tolerance. Cell Host & Microbe 34, 1614–1629.e1618 (2026). 10.1016/j.chom.2026.06.019

81 Kim, D., Paggi, J. M., Park, C., Bennett, C. & Salzberg, S. L. Graph-based genome alignment and genotyping with HISAT2 and HISAT-genotype. Nature Biotechnology 37, 907–915 (2019). 10.1038/s41587-019-0201-4

82 Liao, Y., Smyth, G. K. & Shi, W. featureCounts: an efficient general purpose program for assigning sequence reads to genomic features. Bioinformatics 30, 923–930 (2014). 10.1093/bioinformatics/btt656

83 Love, M. I., Huber, W. & Anders, S. Moderated estimation of fold change and dispersion for RNA-seq data with DESeq2. Genome Biol 15, 550 (2014). 10.1186/s13059-014-0550-8

84 Zhu, A., Ibrahim, J. G. & Love, M. I. Heavy-tailed prior distributions for sequence count data: removing the noise and preserving large differences. Bioinformatics 35, 2084–2092 (2019). 10.1093/bioinformatics/bty895

85 Subramanian, A. et al. Gene set enrichment analysis: a knowledge-based approach for interpreting genome-wide expression profiles. Proceedings of the national academy of sciences 102, 15545–15550 (2005).

86 Liberzon, A. et al. The Molecular Signatures Database (MSigDB) hallmark gene set collection. Cell Syst 1, 417–425 (2015). 10.1016/j.cels.2015.12.004

87 Wickham, H. in ggplot2: Elegant graphics for data analysis 11–31 (Springer, 2016).

88 Huang, K. et al. Autonomous biomedical research with an artificial intelligence agent. Science 393, eadz4351 (2026). doi:10.1126/science.adz4351

