## Supplemental figures for "Mucin promotes pneumococcal colonization via autolysis suppression and tryptophan-mediated immune modulation"

**Figure S1.**

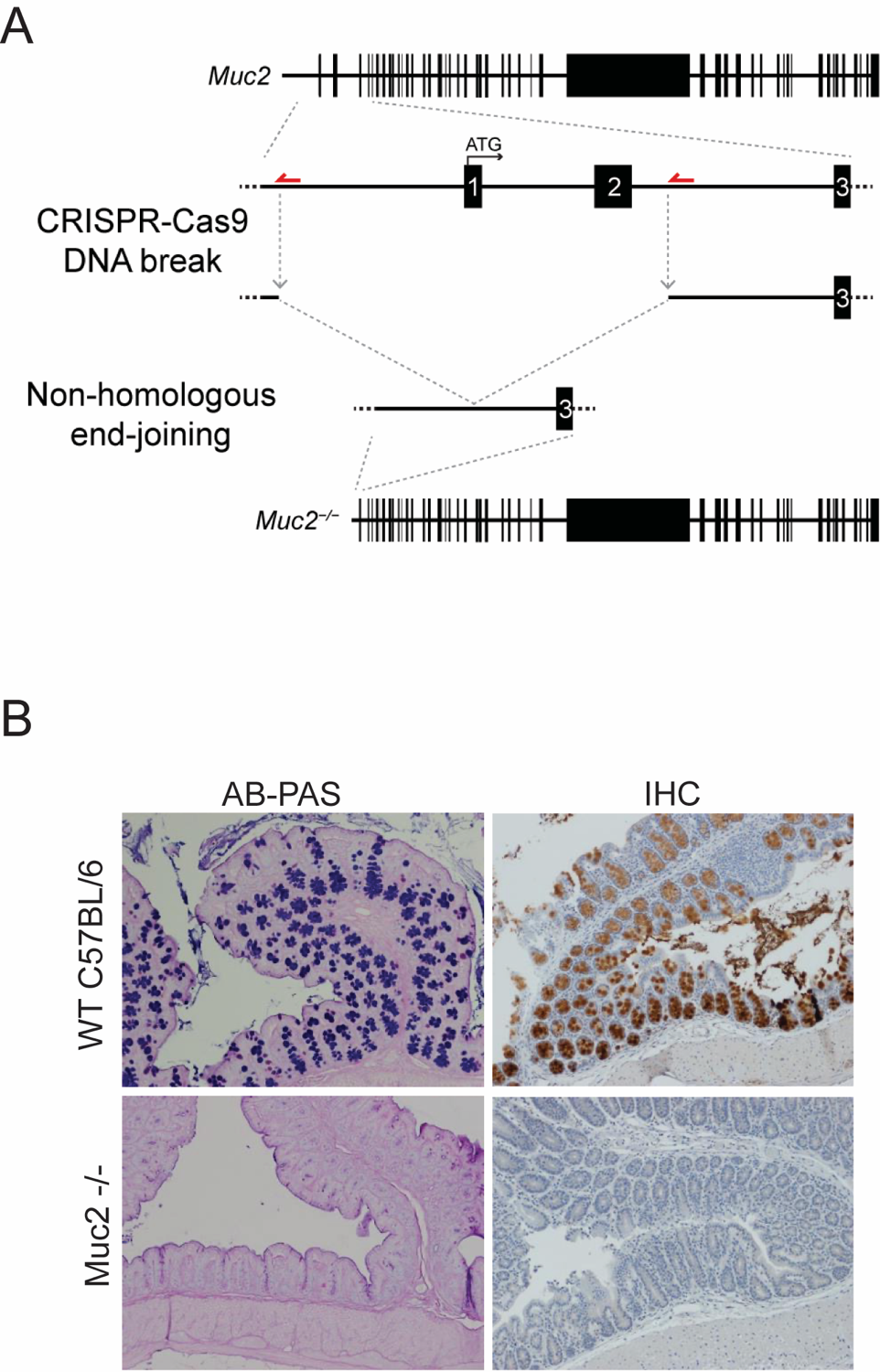

**BHN97 has a shorter colonization period in mice lacking Muc2.** (A) CRISPR strategy for generating *Muc2* mutants in C57BL/6 mice. (B) Muc2 production in intestines of *Muc2* -/- and wild type C57BL/6 mice via antibody staining and IHC.

**Figure S2.**

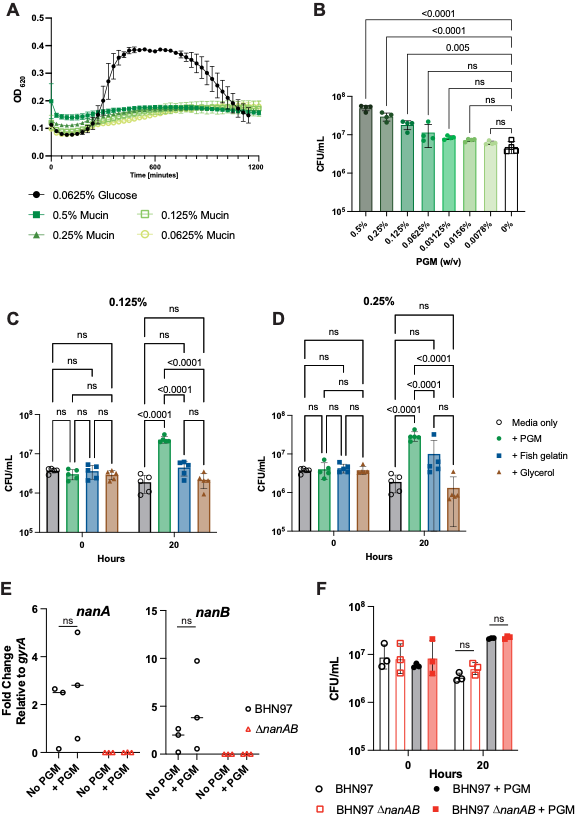

**Prolonged stationary phase is a phenotype specific to PGM.** (A) BHN97 was grown in defined media lacking a carbon source, supplemented with either glucose or PGM. (B) Growth of BHN97 in media containing 0.0625% glucose and various amounts of PGM. (C) Growth of BHN97 in media supplemented with osmoprotectants. (E) qPCR of *nanA* and *nanB* expression in wild type BHN97 and an isogenic ∆*nanAB* double mutant grown with or without mucin. (F) CFU enumeration of wild type and the ∆*nanAB* mutant grown with or without mucin for 20 hours; mutant still exhibits prolonged stationary phase when grown in media supplemeted with PGM.

**Figure S3.**

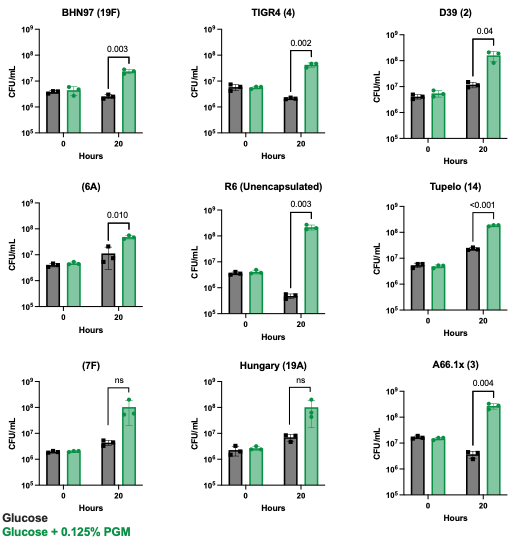

**PGM supplementation across multiple pneumococcal isolates.** Strain names and capsule types, in parenthesis, listed above each graph. All replicates performed in biological triplicate. Recoverable CFU from 4 mL cultures grown for 20 hours in designated media. Significance determined by unpaired t tests; non-significant values are not displayed.

**Figure S4.**

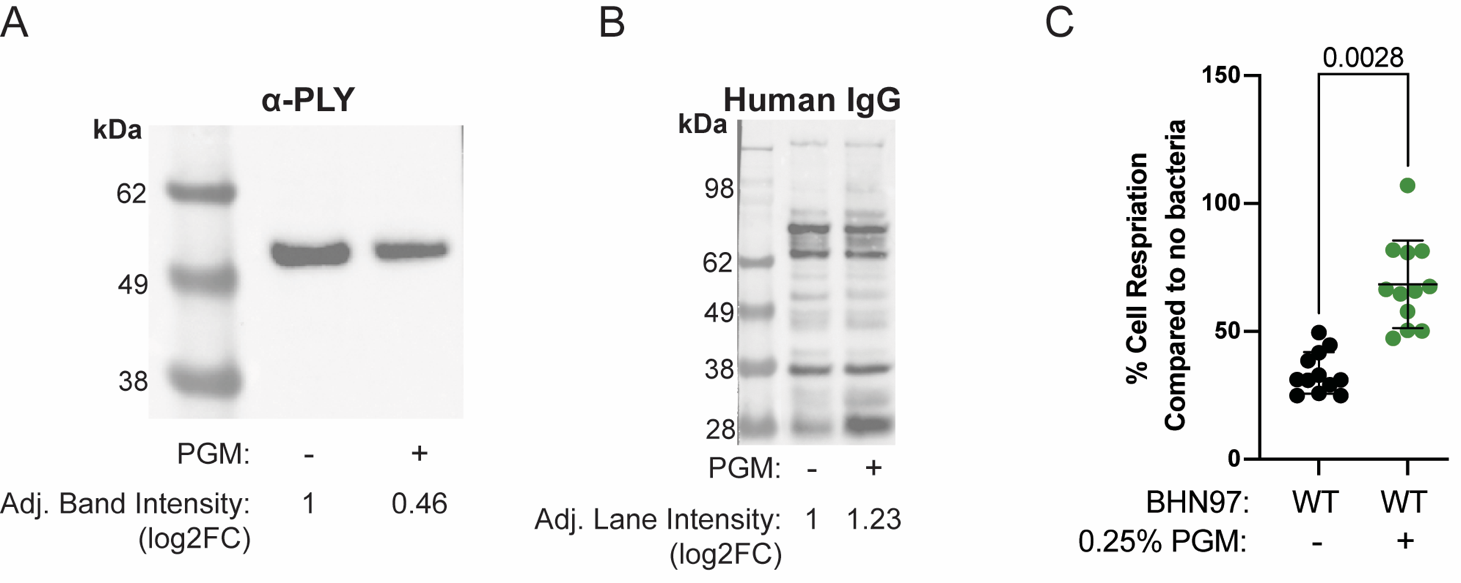

**Pneumococci grown in PGM release less pneumolysin into the supernatant.** (A) Western blot of pneumolysin (PLY) of cells grown with or without PGM. (B) Loading control for (A), showing all proteins reactive with human IgG. Intensity determined with Bio-Rad Image lab Software. (C) AlamarBlue assay on A549 cells incubated with pneumococcal spent media for 1 hour to measure cell respiration. Mean and SD graphed of biological quadruplicates performed in technical triplicate. Significance determined by one-way ANOVA.

**Figure S5.**

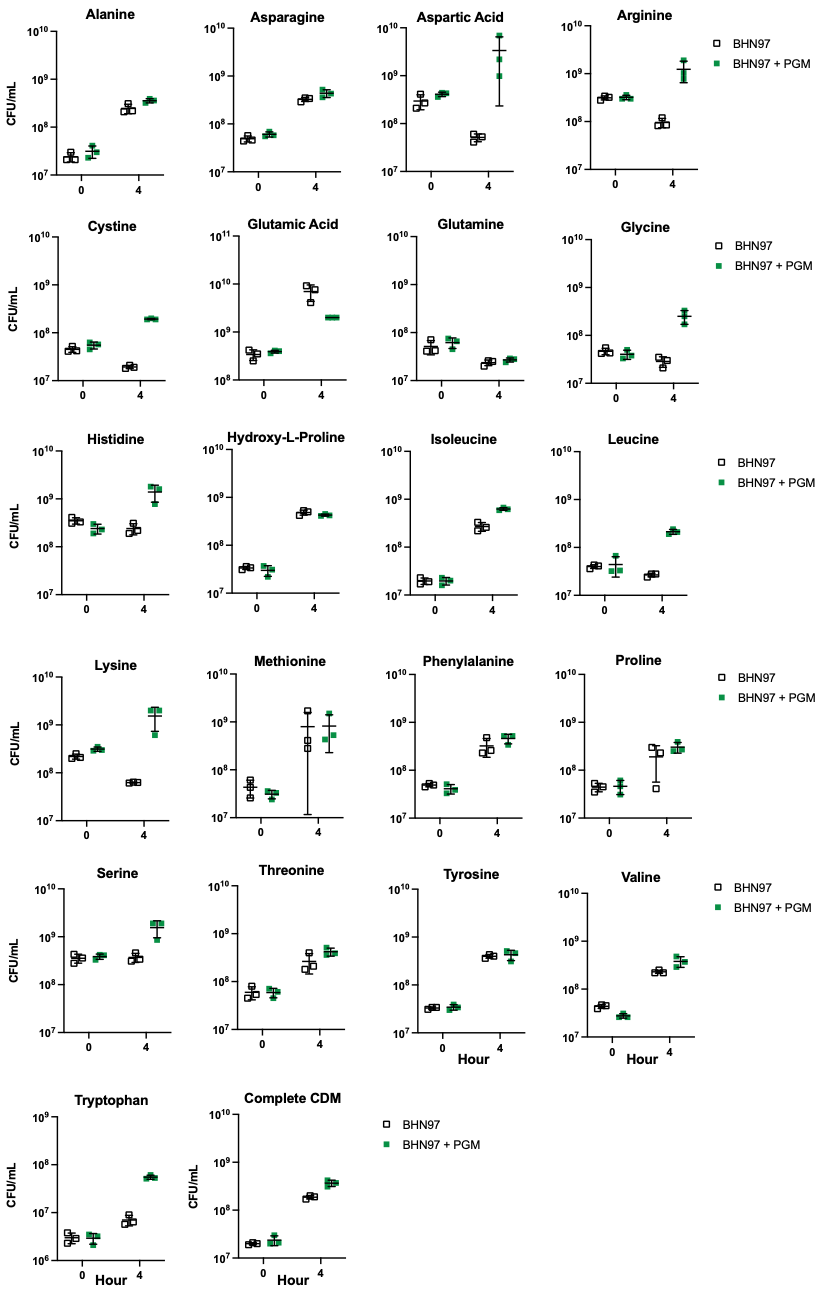

**Mucin rescues pneumococcal growth defects in amino acid limited conditions.** BHN97 was grown in CDM lacking individual amino acids, with or without PGM supplementation, in biological triplicate.

**Figure S6.**

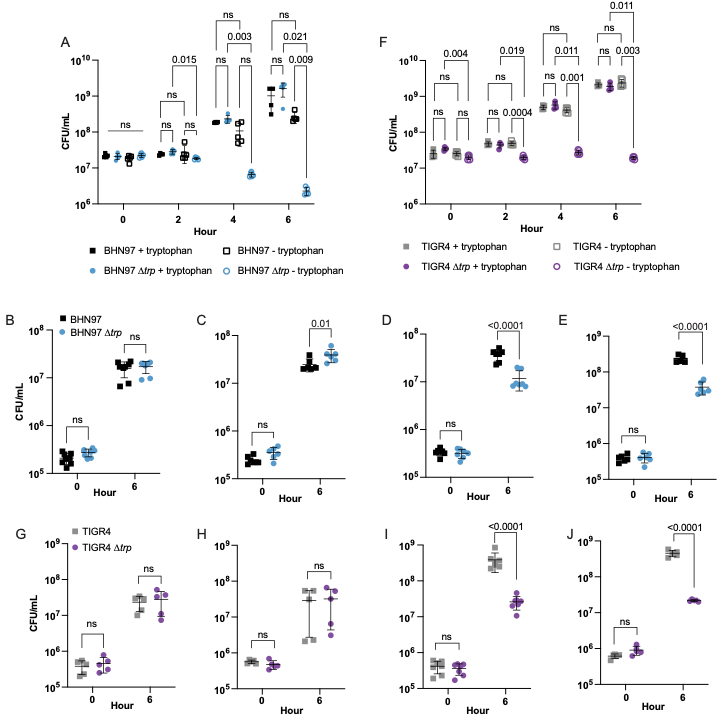

**Pneumococci lacking the tryptophan biosynthetic operon have a growth defect in whole blood.** ∆*trp* mutants in BHN97 (A) and TIGR4 (F) cannot grow in the absence of tryptophan. 5 biological replicates, significance determined by 2-way ANOVA with Tukey’s multiple comparisons. BHN97 ∆*trp* grows comparably to wild type in mouse (B) and human (C) serum, but not mouse (D) or human (E) whole blood. TIGR4 ∆*trp* grows comparably to wild type in mouse (G) and human (H) serum, but not mouse (I) or human (J) whole blood. N = 5 to 7 biological replicates. Significance determined by 2-way ANOVA with Sidak’s multiple comparisons.

**Figure S7.**

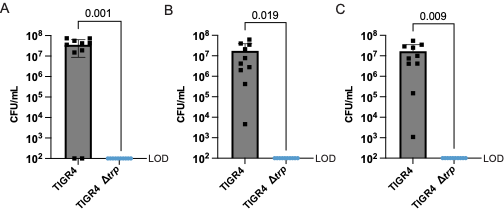

**TIGR4 ∆*trp* cannot cause invasive disease.** Mice were infected intraperitoneally with wild type or ∆*trp* TIGR4. 21 hours post infection, blood (A), spleen (B), and kidneys (C) were harvested, and pneumococci were enumerated. Significance determined by Mann-Whitney test.
